# Selective molecular inhibition of the HDAC6 ZnF-UBP binding domain impairs multiple myeloma cell proliferation

**DOI:** 10.1101/2024.05.14.594071

**Authors:** Rafael Rincón, Isabel F. Coira, Antoine Richieu, Fedaa Attana, Muriel Urwyler, Shozeb Haider, Carole Bourquin, Philippe Bertrand, Muriel Cuendet

## Abstract

Multiple myeloma is a plasma cell malignancy with poor prognosis despite the recent development of new therapeutic options. Histone deacetylase 6 (HDAC6) is overexpressed in multiple myeloma patients and may be involved in the acquisition of resistance to conventional anti-proteasome treatments. Beyond displaying a deacetylase catalytic activity, HDAC6 can recognize ubiquitinated motifs from misfolded proteins through its C-terminal ZnF-UBP binding domain and send the defective proteins to the aggresome for degradation. Here, we explore the role of the ZnF-UBP binding domain of HDAC6 in the function of multiple myeloma cells. A non-functional ZnF-UBP domain containing a 2-residue mutation in the binding site was designed and the absence of ubiquitin binding was confirmed in a cell-free assay. Molecular docking simulations and electrostatic calculations revealed a significant decrease in the electrostatic potential of the mutated peptide, which is crucial for the stability of the complex with ubiquitin. A multiple myeloma cell line containing the non-functional ZnF-UBP domain was then engineered. Although the deacetylase activity of HDAC6 was maintained in these cells, they showed reduced cell growth, impaired aggresome formation and a dysregulated gene expression profile that was more pronounced than cells entirely deficient in HDAC6. These results indicate that a non-functional ZnF-UBP binding domain impacts the function of multiple myeloma cells. Based on these findings, a series of quinazolinylpropanoic acid derivatives was synthesized to explore the inhibitory activity of small molecules to this domain. We propose that ZnF-UBP binding domain inhibitors should be further evaluated as potential therapeutic agents in multiple myeloma.

## 1. Introduction

Multiple myeloma (MM) is an incurable plasma cell malignancy accounting for about 10% of hematological malignancies. The uncontrolled growth of monoclonal plasma cells in the bone marrow leads to complications such as anemia and destructive bone lesions, followed by hypercalcemia and renal failure (1). Considered a late-onset disease, MM has a median age of about 70 years at diagnosis and a global incidence of 4.5 – 6 per 100,000 per year, which is continuously increasing due to population growth and aging (2). During the last decades, overall survival and quality of life of MM patients has notably improved following the introduction of proteasome inhibitors (e.g., bortezomib, ixazomib and carfilzomib), monoclonal antibodies against MM cell surface antigens (e.g., daratumumab and elotuzumab), immunomodulatory agents (e.g., thalidomide, lenalidomide and pomalidomide) and autologous hematopoietic stem cell transplantation, when applicable (3). However, nearly all patients relapse and become refractory to treatment after a few years of initial therapy, and treatment regimens have to be adapted (4).

A few years ago, histone deacetylase (HDAC) inhibitors emerged as a possible new therapeutic strategy against MM. HDACs are essential epigenetic regulators in the cell and they are involved in the maintenance of gene expression, chromatin stability and cell homeostasis. Dysregulation of HDACs is involved in a large variety of cancers affecting crucial processes such as cell proliferation, cell differentiation, angiogenesis, autophagy and apoptosis (5). Previous studies have shown that some HDACs are overexpressed in MM cell lines, MM primary cells from patients and MM bortezomib-refractory models when compared to their normal plasma cell counterparts, and that they are indicators of poor prognosis (6, 7). The pan-HDAC inhibitor panobinostat showed promising overall response rates in combination with bortezomib and dexamethasone for bortezomib-refractory MM patients in the PANORAMA 1 clinical trial, which led to its FDA accelerated approval in 2015 (8). However, panobinostat treatment leads to important adverse events, such as diarrhea, fatigue, thrombocytopenia and anemia and it was withdrawn from the market in 2022 (9). Isoform-selective inhibition of HDAC6 has been proposed as a therapeutic strategy against MM with a better safety profile than pan-HDAC inhibitors. HDAC6 acts mainly in the cytoplasm, due to its nuclear export signal (NES) and its Ser-Glu tetradecapeptide repeat domain (SE14) for cytoplasmic stabilization (10). It contains two catalytic domains, CD1 and CD2, which deacetylate client proteins such as α-tubulin and Hsp90, among others (11). Its cytoplasmic deacetylase role is thought to be involved in important processes such as microtubule regulation and chaperone-induced stress response (12). In addition, HDAC6 contains a C-terminal zinc finger ubiquitin-binding protein (ZnF-UBP) domain, which can bind to ubiquitinated motifs. Therefore, HDAC6 can recognize the unanchored C-terminal diglycine motifs of ubiquitin that are generated in the protein aggregates by deubiquitinase ataxin-3 (13). This leads to the transport of ubiquitinated misfolded proteins via the microtubule network to the aggresome for further degradation. Given this, HDAC6 has been proposed as a main actor for aggresome formation and autophagy activation, which are necessary for the elimination of cytotoxic protein aggregates (14). This mechanism of protein clearance is thought to be responsible for the resistance to proteasome inhibitors in MM patients, and preclinical studies have shown that HDAC6 inhibitors induced synergistic anti-tumor activity with bortezomib in bortezomib-resistant cell models (15, 16). Moreover, phase I/II clinical trials showed that the HDAC6 inhibitor ricolinostat was better tolerated than panobinostat and that its combination with bortezomib displayed synergistic effects (17, 18). However, it still caused noticeable adverse events that could be explained by the destabilization of the microtubule network. Like other HDAC inhibitors, ricolinostat is an hydroxamate-based small molecule designed to bind the catalytic CD2 domain of HDAC6 and, therefore, it does not target the ZnF-UBP binding domain (19). The available X-ray crystallographic structures of the HDAC6 ZnF-UBP binding domain prompted researchers to design inhibitors that could mimic the terminal glycine-glycine repeat of ubiquitin (GG) that binds to the ZnF-UBP cavity. In their work, Ferreira de Freitas et al. produced a series of propanoic acids bearing various aryle or biaryle groups, with one of the best compounds being a quinazolinylpropanoic acid (20). This compound was active at the micromolar level and X-ray crystallographic data (PDB number: 6CED) revealed how the propanoic chain mimicked the ubiquitin GG repeat in the binding domain. More recently, the same group achieved the synthesis of another inhibitor based on the same scaffold, and X-ray data (PDB number: 8G45) showed that a conveniently selected N_3_-substituent could improve the docking capability (21).

In the current study, the molecular and chemical inhibition of the HDAC6 ZnF-UBP binding domain was explored in a MM cell line model to understand the role this domain plays in MM pathogenesis. Its specific inhibition may provide a rationale for a new approach to block the aggresome pathway – alone or in combination with proteasome inhibitors – that may limit the side effects that pan-HDAC or HDAC6 inhibitors induce in patients.

## 2. Results

### 2.1 R1155A-Y1156A mutation in the HDAC6 ZnF-UBP binding domain impaired its interaction with ubiquitin in a cell-free assay

A two-residue mutation was proposed to inhibit the binding of ubiquitin to the HDAC6 ZnF-UBP domain. Previous crystallography data of the human HDAC6 ZnF-UBP binding domain bound to the ubiquitin C-terminal pentapeptide RLRGG showed that residues R1155 and Y1156 were essential to anchor these last amino acids of ubiquitin to the ZnF-UBP cavity (22). The polar side chains of R1155 and Y1156 were oriented towards the inside of the cavity and established hydrogen bonds with RLRGG (Fig. 1). Therefore, a ZnF-UBP peptide (HDAC6^1109 – 1215^), in which residues R1155 and Y1156 were transformed into alanines, was generated through site-directed mutagenesis and peptide purification. The original backbone of the peptide was maintained and the polar side chains where RLRGG pentapeptide was able to interact were removed. Peptide expression was confirmed through SDS-polyacrylamide gel electrophoresis (SDS-PAGE). The binding affinity of the wild-type (ZnF-UBP^WT^) and mutated (ZnF-UBP^RY^) peptides to the RLRGG FITC-labeled ubiquitin pentapeptide was assessed using a fluorescence polarization assay. The RLRGG pentapeptide was incubated at 50 nM with increasing concentrations of the ZnF-UBP^WT^ and ZnF-UBP^RY^ peptides, and results showed that only the ZnF-UBP^WT^ peptide could interact with RLRGG (Fig. 2). ZnF-UBP^RY^ did not interact with the pentapeptide, suggesting that the residues R1155 and Y1156 of HDAC6 are essential for its ubiquitin-binding capacity.

**Figure 1.**
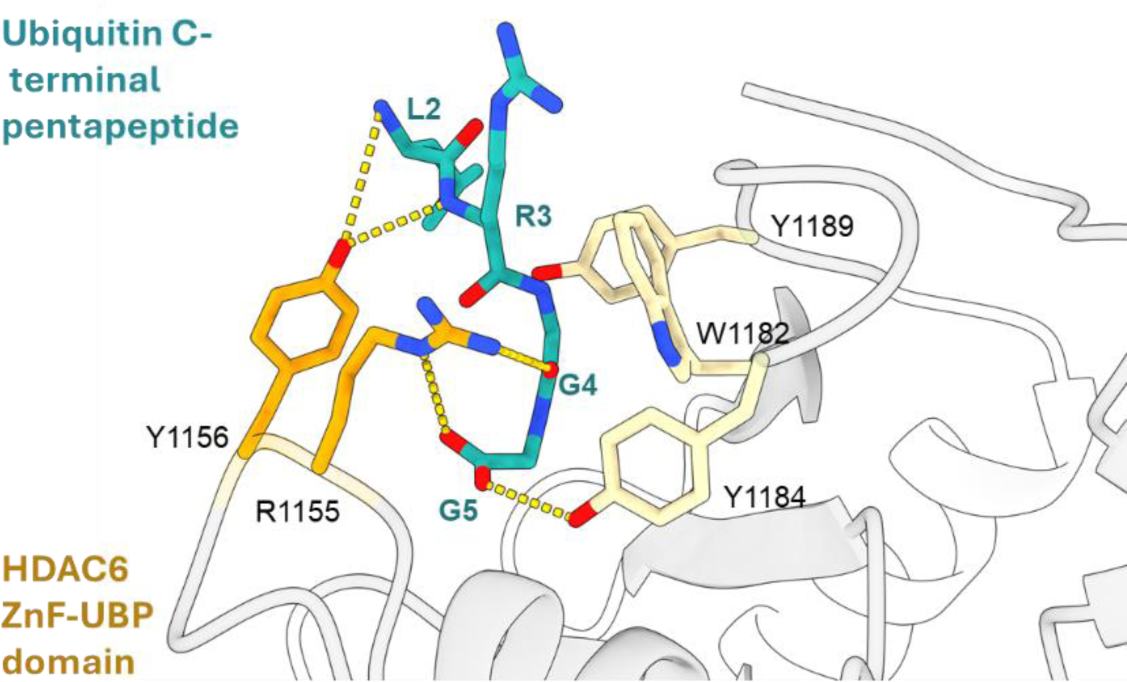
Representation of the human HDAC6 ZnF-UBP binding domain bound to the ubiquitin C-terminal peptide RLRGG. Guanidine and aromatic side chains of the residues R1155 and Y1156 establish hydrogen bonds to anchor the last four amino acids of ubiquitin.

**Figure 2.**
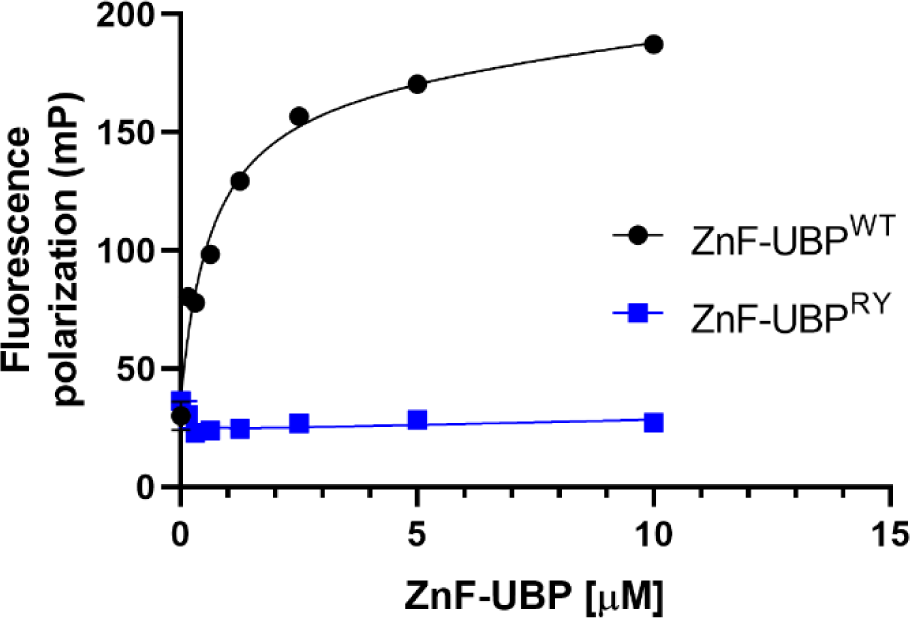
R1155A-Y1156A mutations in the HDAC6 ZnF-UBP domain impaired the interaction with ubiquitin. Fluorescence polarization (FP) saturation curves using increasing concentrations of ZnF-UBP^WT^ and ZnF-UBP^RY^ peptides ZnF-UBP domain with a constant concentration of the FITC-labeled RLRGG pentapeptide (50 nM). A k_d_ of 0.68 ± 0.14 μM was obtained from the average of three independent experiments for the ZnF-UBP^WT^ peptide.

### 2.2. R1155A-Y1156A mutation in the HDAC6 ZnF-UBP binding domain altered its surface electrostatic charge potential

To rationalize the structural role of the R1155A-Y1156A mutation, molecular dynamics simulations of the ZnF-UBP^WT^ peptide (apo), in complex with RLRGG and the ZnF-UBP^RY^ peptide were carried out. The backbone conformation of the apo ZnF-UBP^WT^ and the RLRGG pentapeptide complex is well conserved in all simulations. The side chains of residues R1155 and Y1156 are highly dynamic and act as gatekeepers to the ubiquitin binding site. This is consistent with previously reported studies (13, 20). The principal components analysis identified that mutating R1155 and Y1156 side chains have negligible effect on the overall dynamics of the ubiquitin binding site (Fig. 3A). This observation did not explain why the ubiquitin pentapeptide was unable to bind to the ZnF-UBP^RY^ peptide. Peptide impairment was then investigated by running Poisson-Boltzmann solvation calculations. These calculations revealed a significant decrease in the electrostatic potential, suggesting a substantial reduction in charged and polar interactions that are crucial for the stability of the ZnF-UBP^WT^-RLRGG complex. The change can be visually appreciated in Fig 3B, which highlights the electrostatic charged surfaces of the ubiquitin pentapeptide and the complementary charged surfaces of the ZnF-UBP^WT^ and the ZnF-UBP^RY^ mutant. A clear diminished electrostatic charge complementarity between the pentapeptide and the ZnF-UBP^RY^ mutant was observed. Furthermore, the R1155A-Y1156A mutation also led to the loss of hydrogen bonds between the mutated side chains and the backbone/side chain of the pentapeptide, further undermining the structural integrity of the interaction. Following the analysis of the atomistic interactions between the pentapeptide and ZnF-UBP showed an altered contact frequency of residues of the pentapeptide when comparing the wild-type and the mutated forms (Fig 3C).

**Figure 3.**
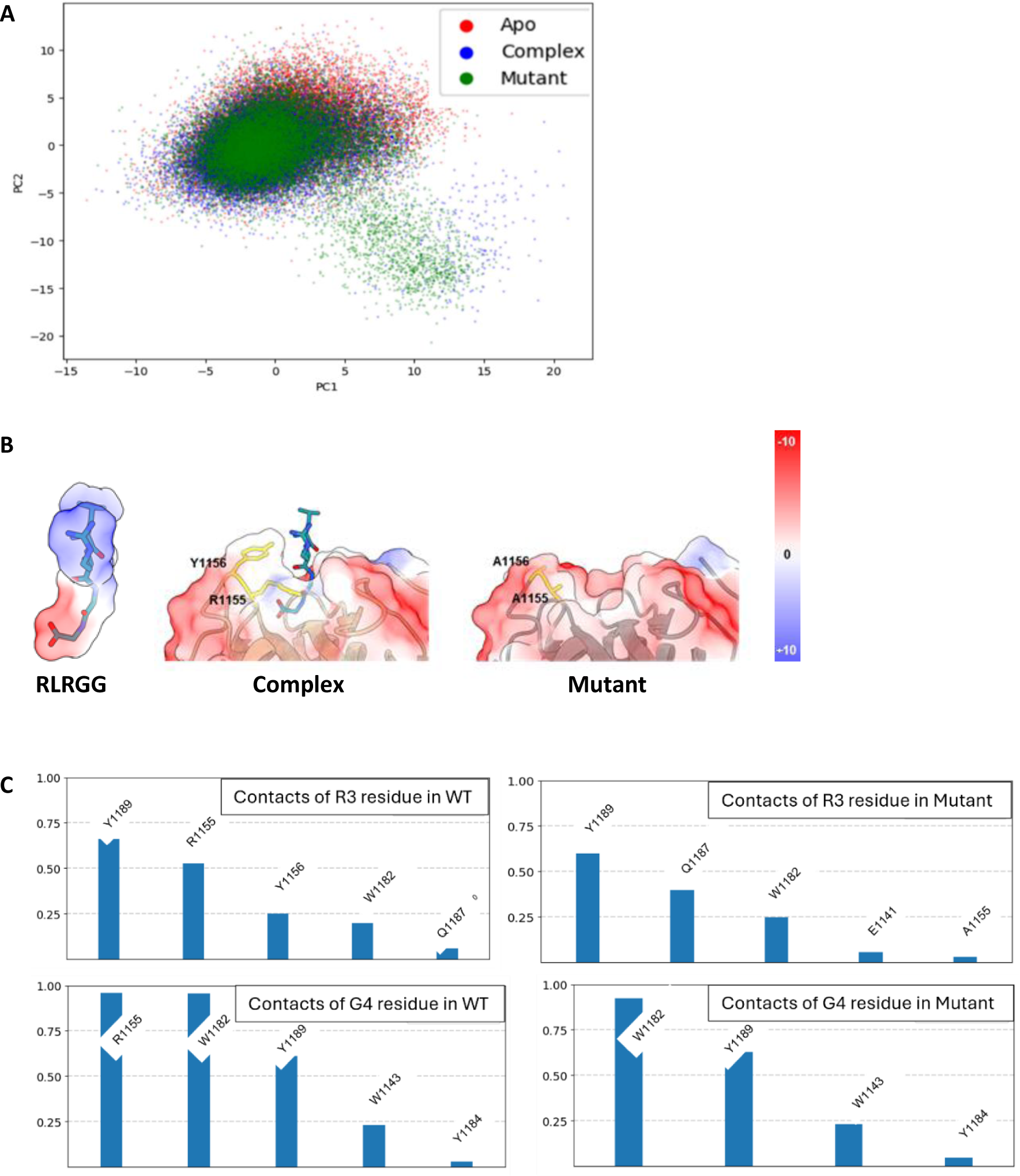
Some sentence. (**A**) Principal component analysis projection of the backbone positions for the ZnF-UBP^WT^ peptide (Apo), in complex with RLRGG (Complex) and the ZnF-UBP^RY^ peptide (Mutant). (**B**) Representation of the calculated electrostatic charged surface of the ubiquitin terminal pentapeptide RLRGG, the ZnF-UBP^WT^ peptide in complex with RLRGG (Complex) and the ZnF-UBP^RY^ peptide (Mutant). (**C**) Bar charts highlighting contact frequencies of R3 and G4 residues of the ubiquitin C-terminal pentapeptide calculated at 3.5 Å with nearest 4 bonded residues excluded.

### 2.3. Molecular inhibition of the HDAC6 ZnF-UBP binding domain did not impair HDAC6 expression or its catalytic activity but impaired aggresome formation in RPMI 8226 cells

The evidence of ZnF-UBP molecular inhibition through a two-residue mutation was the rationale for generating a MM cell line with a non-functional HDAC6 ZnF-UBP binding domain. Two RPMI 8226 cell lines were generated through CRISPR-Cas9 technology: a full HDAC6 knockout cell line (HDAC6^KO^) and a ZnF-UBP-mutated cell line (HDAC6^RY^). After delivery of the Cas9 ribonucleoprotein (RNP) complex through electroporation and subsequent clonal expansion, the efficiency of the CRISPR-Cas9 edition was verified at the DNA level for clone selection. Tracking of indels by decomposition (TIDE) analysis showed that the HDAC6^KO^ RPMI 8226 cells successfully acquired indels of various lengths in exon 3, a modification that is normally deleterious for protein expression due to reading frame shifting. Sequencing of the genomic region encoding for ZnF-UBP binding domain showed that reference codons CGT TAC (R1155, Y1156) were mutated into GCT GCC (A1155, A1156) in HDAC6^RY^ RPMI 8226 cells.

At the protein level, HDAC6^KO^ cells did not express HDAC6 and consistently, acetylated tubulin expression increased with the loss of HDAC6, as it is one of its main substrates (23). HDAC6^RY^ cells maintained HDAC6 expression despite the change in residues R1155 and Y1156 into nonpolar alanines. No changes in tubulin acetylation were observed, suggesting that this two-residue mutation in the ZnF-UBP binding domain did not affect HDAC6 expression or tubulin deacetylase activity (Fig. 4A). Total HDAC catalytic activity was also measured using a UHPLC-MS method in which the ratio between the deacetylated and acetylated MAL, a pan-HDAC substrate, was measured. The total HDAC catalytic activity was maintained in HDAC6^RY^ cells compared to the wild-type cells, while a reduction in the deacetylation of MAL substrate was observed in HDAC6^KO^, which is consistent with the loss of HDAC6 expression (Fig. 4B). The aggresome formation was then measured in the different RPMI 8226 cell lines to assess the role that HDAC6 plays in this process. Cells were stimulated with the proteasome inhibitor MG-132 (5 µM) for 18 h to induce aggresome formation in response to proteasome blockade. A decrease in aggresome activation was observed in HDAC6^KO^ and HDAC6^RY^ cells compared to wild-type cells (Fig. 4C), indicating that a functional HDAC6 ZnF-UBP binding domain may be necessary for a correct aggresome formation.

**Figure 4.**
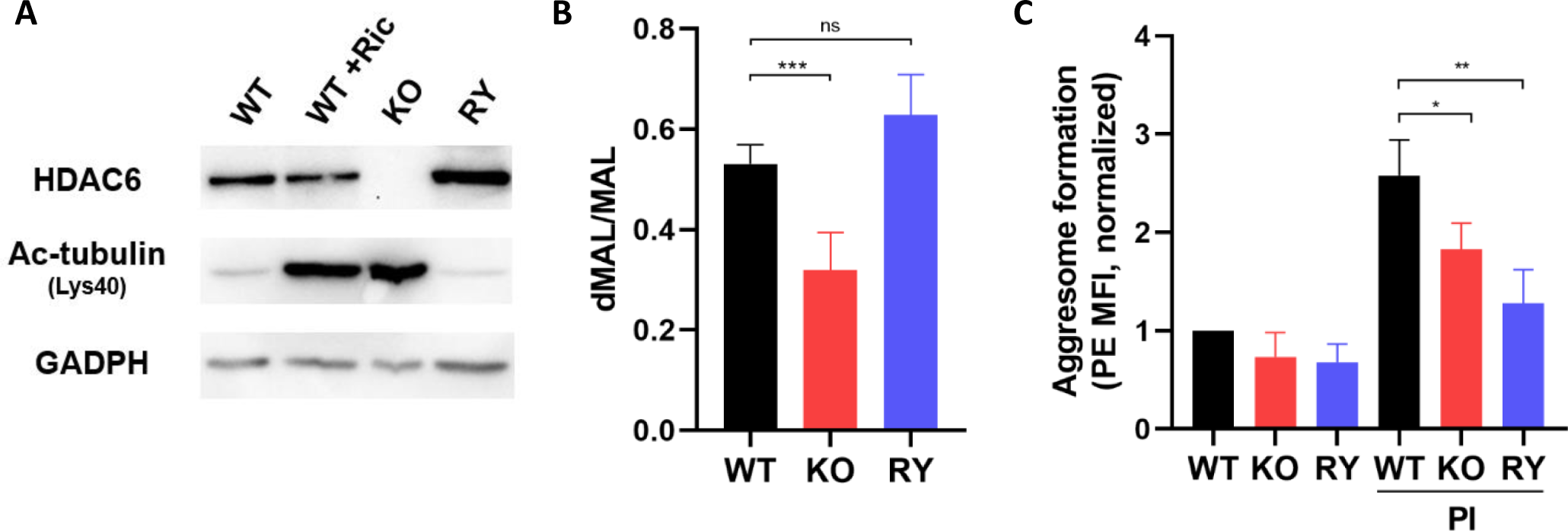
The molecular inhibition of the HDAC6 ZnF-UBP binding domain did not impair the HDAC6 expression or its catalytic activity in RPMI 8226 cells but impaired the aggresome formation. (**A**) Expression of HDAC6 and acetylated tubulin in wild-type, HDAC6^KO^ and HDAC6^RY^ RPMI 8226 cell lines (WT, KO, RY) as determined by western blotting. Wild-type RPMI 8226 cells treated with 2 µM ricolinostat (WT + Ric) for 24 h were used as positive control. (**B**) Deacetylase activity of the three RPMI 8226 cell lines (WT, KO, RY) after incubation with the MAL substrate was obtained by measuring the ratio deacetylated MAL (dMAL)/MAL by UHPLC-MS. Results are the average of three independent experiments. (**C**) Aggresome activation was induced through a proteasome inhibitor (PI: MG-132, 5 µM) for 18 h and measured by flow cytometry (PE mean fluorescence of intensity). Results are the average of three independent experiments. Groups were compared using unpaired t test, ns: non-significant, * p < 0.05, ** p < 0.01, *** p < 0.001.

### 2.4. Proliferation of HDAC6^KO^ and HDAC6^RY^ RPMI 8226 cells was reduced and cell cycle was dysregulated compared to wild-type cells

Cell growth and cell cycle were assessed in the different RPMI 8226 cell lines to determine whether the loss or mutation of HDAC6 could impact cell proliferation. Cell growth was decreased in HDAC6^KO^ and HDAC6^RY^ cells compared to wild-type cells with statistically significant differences after 72 h (Fig. 5A). In agreement with these findings, significant changes in the cell cycle were observed, with a decrease of cells in the G0/G1 phase and an increase of cells in the S and G2/M phases in HDAC6^KO^ and HDAC6^RY^ cells compared to wild-type cells (Fig. 5B). These results suggest that factors such as cell cycle arrest or extended cell cycle phase durations may affect the progression of the cell cycle in HDAC6^KO^ and HDAC6^RY^ cells.

**Figure 5.**
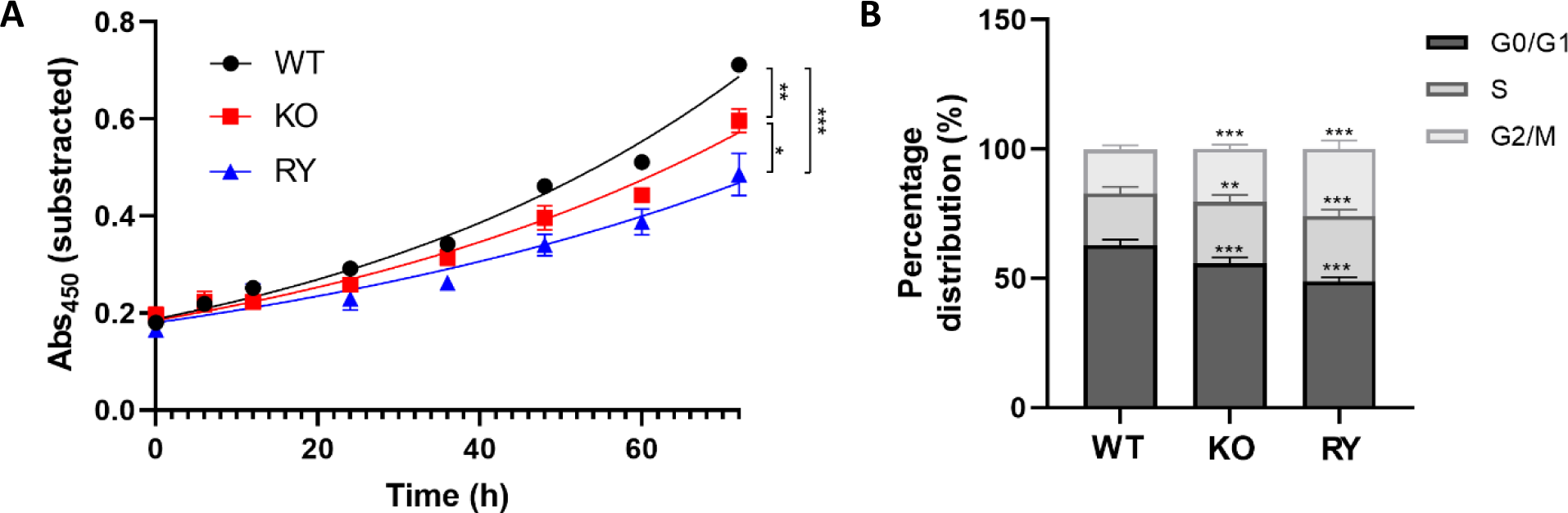
The molecular inhibition of the HDAC6 ZnF-UBP binding domain and the full knockout of HDAC6 decreased cell proliferation and deregulated the cell cycle. (**A**) Cell growth was measured in the three RPMI 8226 cell lines (WT, KO, RY) by measuring the metabolic activity. (**B**) Cell cycle distribution in the three RPMI 8226 cell lines (WT, KO, RY) was measured by flow cytometry (propidium iodide). Results are the average of three independent experiments. Groups were compared using unpaired t test, * p < 0.05, ** p < 0.01, *** p < 0.001.

### 2.5. Chemical synthesis of quinazolinones **1a-g** as potential HDAC6 ZnF-UBP inhibitors

Results obtained with the engineered cell lines suggested that a specific inhibition of the HDAC6 ZnF-UBP binding domain might be a valid target for MM therapy. Therefore, selective ZnF-UBP inhibitors with no affinity for the catalytic domains were designed and synthetized. Alternative substitutions of the quinazolinone scaffold were developed, focusing on the benzene part of the quinazolinone. The synthesis of the substituted quinazolinones **1a-g** was based on the procedure published by Ferreira de Freitas et al. for unsubstituted quinazolinone (**1a**, R=H) (Scheme 1) (20). The preparation of compounds **4a-g** was optimized compared to the previous two-step procedure with a more sustainable single step preparation under micro waves conditions using pinane as solvent (24). Then, an esterification led to compounds **5a-g** and the subsequent N-methylation afforded compounds **6a-g** in variable yields. In some cases, an alternative approach was found more productive by performing in one step the esterification as methyl ester and the N-methylation using 3 eq. of CH_3_I and K_2_CO_3_. A final ester hydrolysis gave the expected quinazolinylpropanoic acids **1a-g**. All these new compounds were characterized by NMR and exact mass, and all tested compounds were 95% or more pure as determined by HPLC.

**Scheme 1.**
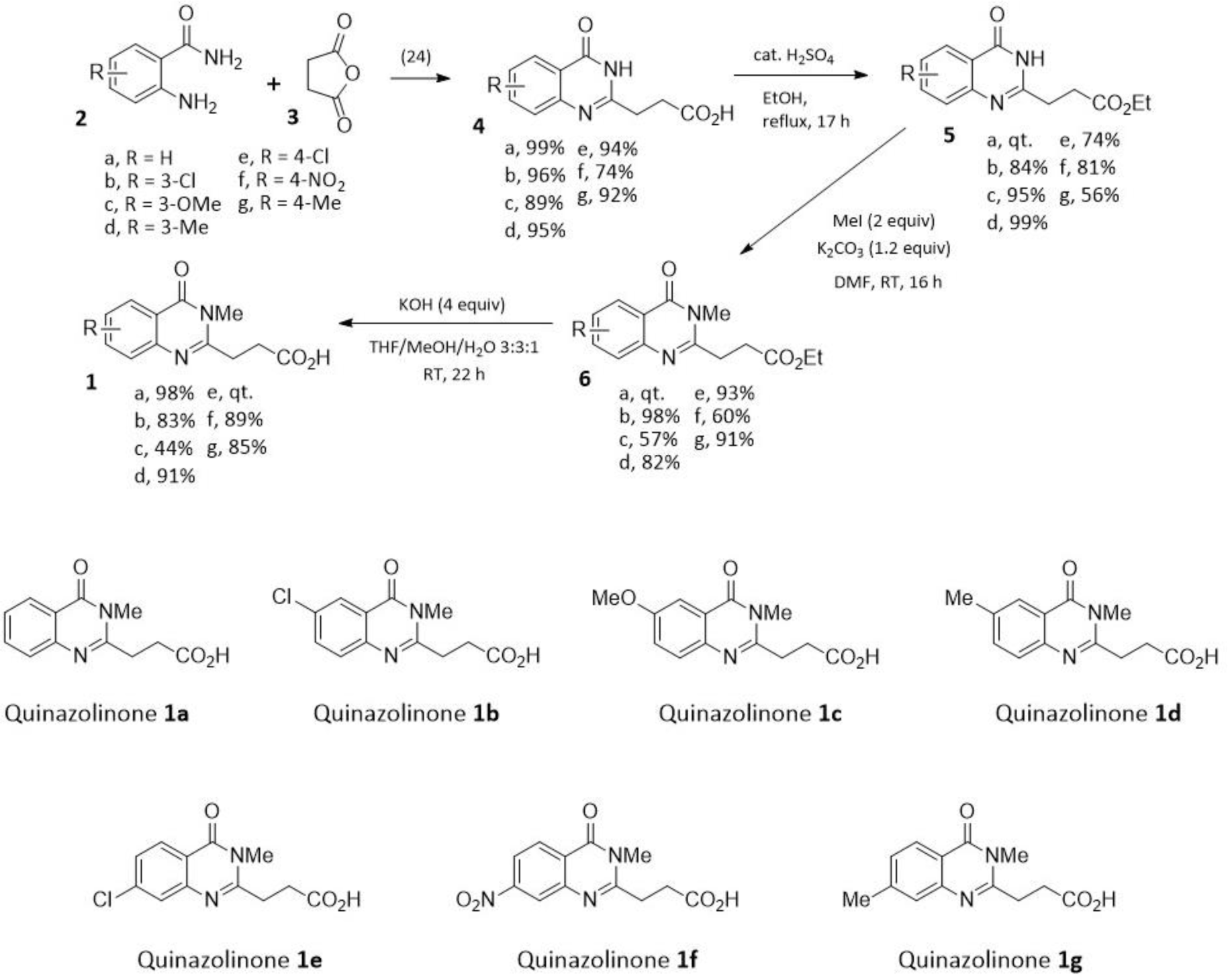
Preparation and structure of quinazolinones **1a-g** as potential ZnF-UBP inhibitors.

### 2.6. Molecular docking of quinazolinones 1a-g

The structural binding of compounds **1a-g** to the HDAC6 ZnF-UBP binding domain were assessed using automated docking. Previous structural studies by Ferreira de Freitas et al. (PDB number: 6CE6) were used as a starting point for chemical exploration around the ZnF-UBP binding site (20). All compounds (**1a-g**) docked in a similar orientation, sandwiched between R1155 and W1182. The pyrimidine ring in the quinazolinone made π-π stacking interactions with W1182 and the carbonyl group was oriented towards Y1189 (Fig. 6). The propanoic acid tail made hydrogen bonding interactions with the side chains of R1155 and Y1184 and the backbone nitrogen of G1154. Compound **1f** was also able to make a hydrogen bond with the side chain of Q1187. The RTCNN scores, used to rank the compounds, ranged between -31.33 and -36.66 (Table 1).

**Figure 6.**
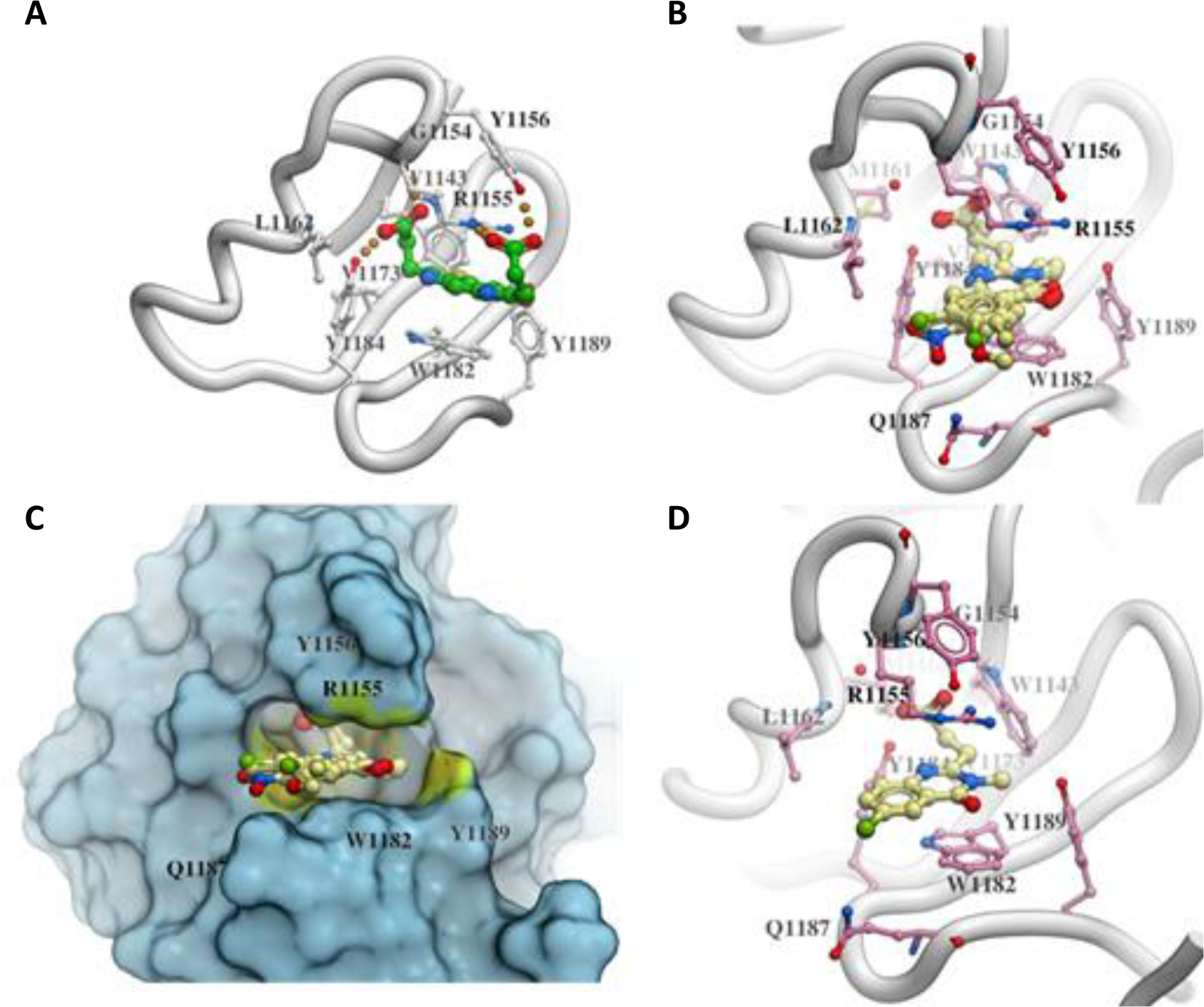
Automated ligand docking. (**A**) Reference compound from the crystal structure 6CE6; (**B**) docked orientation of compounds **1a-g**; (**C**) surface representation of the HDAC6 ZnF-UBP domain with the docked ligands highlighting the binding pocket and (**D**) docked orientation of compound **1b**.

**Table 1.** RTCNN docking scores and inhibition of the HDAC6 ZnF-UBP – RLRGG ubiquitin pentapeptide interaction (IC_50_ in µM) of the ZnF-UBP inhibitor candidates **1a-g**.

| Compounds | RTCNN score<sup>a</sup> | IC<sub>50</sub> [$\mu\text{M}$]<sup>b</sup> |
| --- | --- | --- |
| **1a** | -35.41 | 2.7 ± 0.3 |
| **1b** | -36.66 | 1.9 ± 0.2 |
| **1c** | -31.33 | 2.1 ± 0.5 |
| **1d** | -34.74 | 13.6 ± 1.0 |
| **1e** | -35.89 | 1.9 ± 0.1 |
| **1f** | -32.93 | 2.3 ± 0.2 |
| **1g** | -33.95 | 2.1 ± 0.1 |
| Ricolinostat<sup>c</sup> |  | > 20 |
| Tubacin<sup>c</sup> |  | > 20 |
| Panobinostat<sup>c</sup> |  | > 20 |

<sup>a</sup>The RTCNN score from molecular docking.
<sup>b</sup>Results represent the mean ± standard error of at least three independent experiments performed in triplicate.
<sup>c</sup>The HDAC6 inhibitors ricolinostat, tubacin and panobinostat were used as controls.

### 2.7. ZnF-UBP inhibitor candidates **1a-g** disrupted the HDAC6 ZnF-UBP interaction with ubiquitin in a cell-free assay, but did not show significant effects in RPMI 8226 cells

Compounds **1a-g** were tested for their ubiquitin-displacing activity towards the HDAC6 ZnF-UBP binding domain. Compound **1a** exerted a ubiquitin-displacing activity similar to what was previously published by Ferreira de Freitas et al. (20). Compounds **1b**, **c**, **e** and **g** displayed a slightly improved ubiquitin-displacing activity (Table 1). These results were consistent with RTCNN docking scores, which allowed to identify the strongest binders. The HDAC6 inhibitors ricolinostat, tubacin and panobinostat did not exert any ubiquitin-displacing activity at 20 µM, showing their specificity towards the catalytic domains of HDAC6 and validating this assay as a useful tool to assess the specific ZnF-UBP domain inhibitory capacity of small molecules.

Then, the aggresome formation was measured in RPMI 8226 cells treated with compound **1b**, as it showed the strongest ubiquitin-displacing activity. Cells were treated with **1b** for 24 h with concentrations up to 100 µM prior to adding the proteasome inhibitor (MG-132, 5 µM). No significant differences were observed between the treated and control samples, suggesting that the chemical approach to inhibit the aggresome pathway did not lead to comparable effects than what was obtained with the CRISPR-Cas9-edited HDAC6^RY^ RPMI 8226 cells (data not shown). Moreover, compounds **1a-g** did not exert any anti-proliferative activity or deregulation of the cell cycle in RPMI 8226 cells up to 100 µM (data not shown). Furthermore, a 24 h treatment with **1b** did not induce the expression of cell stress-related proteins (data not shown). Finally, combination treatment of **1b** with bortezomib did not display any synergistic effect in RPMI 8226 cells (data not shown). Taken together, these results indicate that the synthetized compounds were able to bind to the ZnF-UBP domain and displace ubiquitin, but no effect could be identified in RPMI 8226 cells.

### 2.8. Non-functional HDAC6 ZnF-UBP binding led to a decrease in signaling pathways involved in cell adhesion and in the immunological function

To gain insight into pathways affected by the absence of HDAC6 or a non-functional HDAC6 ZnF-UBP binding domain, bulk RNA-seq analysis of the three RPMI 8226 cell lines was performed. Out of 21,394 genes analyzed, 3,498 transcripts (16.4%) corresponded to differentially expressed genes (DEG) in HDAC6^KO^ cells compared to wild-type cells, among which 2,237 (10.5%) were downregulated and 1,261 (5.9%) were upregulated. Greater changes were observed in the HDAC6^RY^ cells, where there were 14,334 (67.0%) DEGs compared to wild-type cells, among which 7,506 (35.1%) were downregulated and 6,828 (31.9%) were upregulated. This shows that the absence of HDAC6 protein led to less changes in gene expression than the two-residue mutation of its ZnF-UBP binding domain (Fig. 7A). Gene ontology (GO) enrichment analysis showed that in both HDAC6^KO^ and HDAC6^RY^ cell lines, pathways linked to cell-cell adhesion and to immune responses were among the most significantly dysregulated processes (Fig. 7B). Interestingly, the number of DEGs involved in these processes was higher in HDAC6^RY^ cells than in HDAC6^KO^ cells, and the fold changes were also globally higher in HDAC6^RY^ cells. For example, the transcripts of the three cell surface proteins desmoglein-2 (*DSG2*), N-cadherin (*CDH2*) and ICAM-1 (*ICAM1*) were significantly down-regulated in the HDAC6^RY^ cell line compared to wild-type cells, but not in the HDAC6^KO^ line. The same trend was observed for light chain immunoglobulin genes (*IGLV2-14*, *IGLC3*, *IGLC2*) and immune receptors and transcription factors (*TLR7*, *TLR4*, *IRF4*) (Fig. 7C).

**Figure 7.**
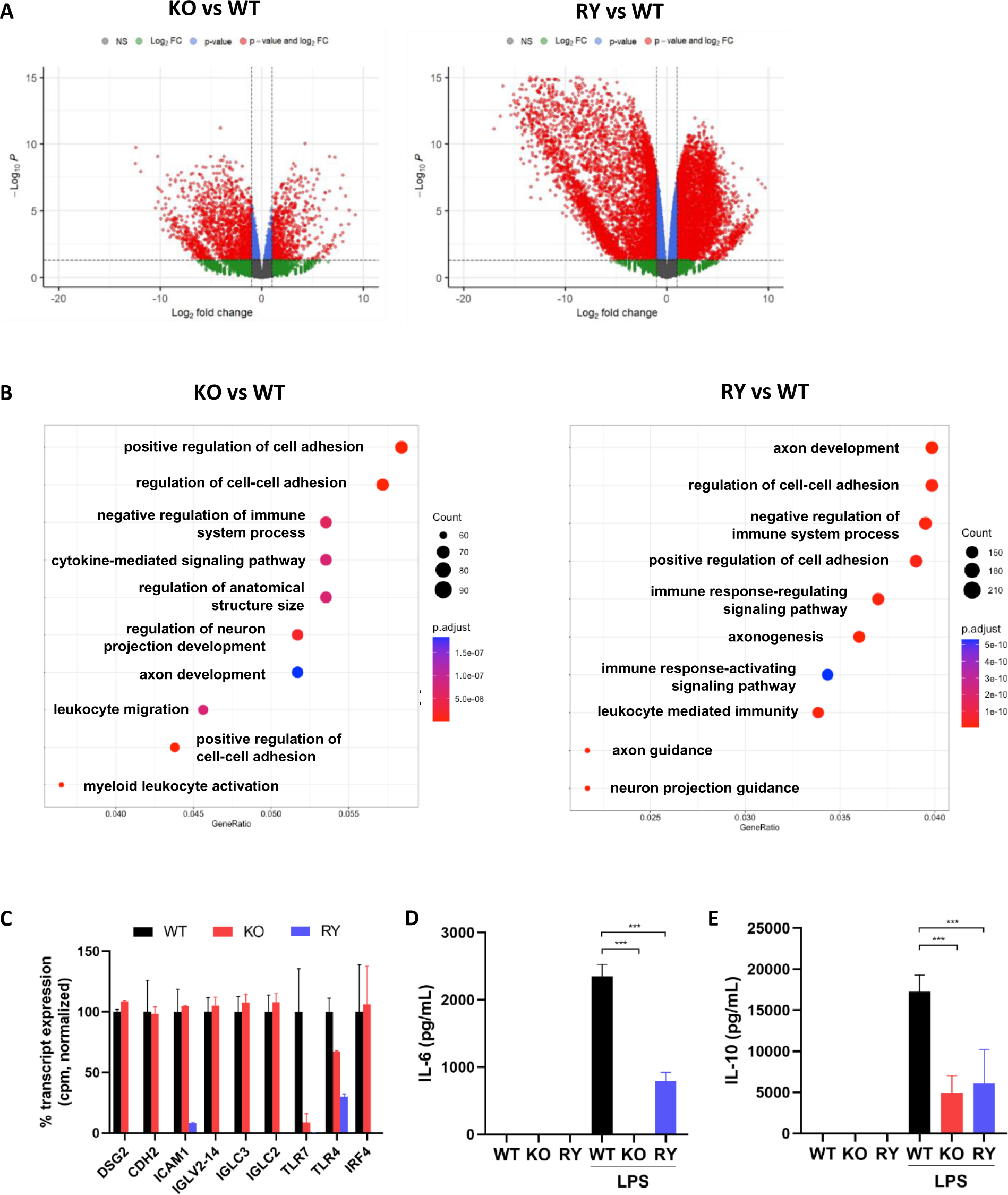
The molecular inhibition of the HDAC6 ZnF-UBP binding domain led to greater transcriptional changes than the full HDAC6 knockout. (**A**) *Volcano* plot of differentially expressed genes (DEGs) in the HDAC6 ^KO^ (KO) and HDAC6^RY^ (RY) RPMI 8226 cell lines compared to wild-type cells, pCutOff = 0.05. (**B**) Gene ontology (GO) enrichment analysis of these DEGs showing the main biological functions significantly enriched in HDAC6^KO^ (KO) and HDAC6^RY^ (RY) RPMI 8226 cell lines compared to wild-type cells. (**C**) Bar plot depicting relative transcript expression of some of these DEGs belonging to cell adhesion pathways, immunoglobulin light chain expression and immune response. (**D**, **E**) IL-6 and IL-10 production stimulated in the three RPMI 8226 cell lines by LPS (100 ng/mL) for 24 h and concentrations measured in the cell supernatant. Results are the average of three independent experiments. Groups were compared using unpaired t test, *** p < 0.001.

To assess whether HDAC6^KO^ and HDAC6^RY^ cells showed differences in immune function compared to wild-type cells, we measured by ELISA the amount of lambda light chain immunoglobulins secreted by the three RPMI 8226 cell lines, but observed no differences between the lines (data not shown). Further, we stimulated all three cell lines with lipopolysaccharide (LPS), a ligand for the innate immune receptor TLR4, and measured the production of the cytokines IL-6 and IL-10 by ELISA. While LPS induced high amounts of IL-6 in wild-type cells, production was significantly decreased in HDAC6^RY^ cells, and it was completely absent in HDAC6^KO^ cells (Fig. 7D). These results suggest that both the catalytic domains and the ZnF-UBP binding domain of HDAC6 may cooperate to support IL-6 production. In contrast, IL-10 production was decreased in a similar manner in HDAC6^KO^ and HDAC6^RY^ cells compared to wild-type cells (Fig. 7E), suggesting that the ZnF-UBP binding domain, rather than the catalytic domains, is involved in the IL-10 inducing function of HDAC6.

## 3. Discussion

HDAC6 inhibition is currently proposed as a possible therapeutic approach for MM. The versatile and essential role that HDAC6 displays in the cell cytoplasm through its multi-domain structure makes it a promising target for drug development pipelines. Despite the extensive studies focusing on its deacetylase activity towards multiple protein substrates, the knowledge about the role that HDAC6 plays in MM progression is still limited.

In this research article, the role of the HDAC6 ZnF-UBP binding domain was explored as a possible target in MM. To do so, the effect of a two-residue mutation in the ZnF-UBP binding domain was shown in a cell-free assay and *in silico* to be a potential disruptor of the HDAC6-ubiquitin interaction. Then, a MM cell line containing this mutation in the ZnF-UBP domain of HDAC6 was developed to obtain insights into the role this domain plays in MM pathogenesis. Impairment of the ubiquitin-binding capacity of the HDAC6 ZnF-UBP domain directly caused a significant decrease in aggresome formation in RPMI 8226 cells, indicating a direct role of HDAC6 in this pathway through this domain. These results bring new evidence on the role that the ZnF-UBP binding domain plays in a cancer cell model. However, partial aggresome activation was still achieved in these cell lines, suggesting that this process might not exclusively depend on HDAC6, but also on other described aggresome activator proteins, such as p62 or USP10 (25, 26).

In addition to aggresome inhibition, a slower cell growth and an increase in the S and G2/M phases of the cell cycle were observed. Anti-cancer drugs such as kinesin-5 inhibitor BRD9876 or Aurora A inhibitor ENMD-2076 have shown anti-proliferative effects in MM cell models and a cell arrest on the G2/M phase (27, 28). These findings suggest that the HDAC6-ubiquitin interaction could also play a role in the regulation of MM cell division and that this inhibition could induce similar effects to the inhibition of mitotic checkpoints. Aggresome formation, cell growth and the cell cycle were surprisingly more affected in cells with a non-functional HDAC6 ZnF-UBP binding domain than in cells with a full HDAC6 knockout. Based on this observation, a specific inhibition of the ZnF-UBP binding domain, allowing HDAC6 catalytic activity to be unaffected, may be a suitable approach against MM.

At the transcriptional level, a gene ontology analysis indicated that the HDAC6 ZnF-UBP domain may be related to the regulation of cell adhesion molecules. Integrins, cadherins, selectins and syndecan-1/CD138 have previously been shown to play a role in the adhesion and communication between MM and bone marrow stromal cells (BMSC) (29). In this regard, some anti-MM treatments designed to target cell adhesion molecules are being developed. Daratumumab, an anti-CD38 antibody, was approved in 2016 by the FDA for MM treatment, as a single agent or in combination with bortezomib or lenalidomide (30). Natalizumab, an anti-integrin subunit α4 antibody, inhibited the adhesion of MM cells to non-cellular and cellular components of the microenvironment and reduced tumor growth, VEGF secretion and angiogenesis in an immunodeficient murine model (31). Therefore, it will be interesting to further study adhesion and migration to characterize the impact that a non-functional ZnF-UBP domain has in MM cells. Moreover, a negative regulation of the immunological function was suggested at the transcriptomic level for HDAC6^KO^ and HDAC6^RY^ cells. At the protein level, the production of IL-10 induced by TLR4 activation was decreased in both HDAC6^KO^ and HDAC6^RY^ cell lines. Previous publications showed that HDAC6 deletion could inhibit IL-10 production in macrophages and in antigen-presenting cells (32, 33). Similarly, LPS-induced IL-6 production was absent with the loss of HDAC6. This finding is in agreement with a previous study, in which ricolinostat induced an anti-inflammatory response in LPS-stimulated macrophages through the suppression of the TLR4-MAPK/NF-κB pathways, which are essential for the induction of IL-6 (34). Although the association between HDAC6 and immunological functions was already described in myeloid cells, this is to our knowledge the first time it is shown in MM cells. IL-6 reduction may be of clinical value, as IL-6 was reported to play a central role in MM progression by mediating the communication between MM cells, osteoblasts and stromal cells in the bone marrow (35). Interestingly, the production of IL-6 was reduced in the HDAC6^RY^ cells, but to a lesser extent than in HDAC6-deficient cells. This suggests an additive activity between the ZnF-UBP binding domain and the catalytic domains of HDAC6 for the induction of IL-6. Taken together, these findings demonstrate that the catalytic and ZnF-UBP domains of HDAC6 may have their own independent functions for certain cellular processes – such as tubulin deacetylation or aggresome formation –, but they may also cooperate for others, such as those involved in immune responses.

The strong impact of a non-functional HDAC6 ZnF-UBP binding domain on several biological processes in MM cells highlights its potential as a drug target. Substituted quinazolinones were synthetized to mimic the ubiquitin glycine-glycine motif bound to the ZnF-UBP domain. Despite their ubiquitin-displacing activity and their predicted binding *in silico*, the lack of aggresome inhibitory activity of these first designed quinazolinones showed that the cellular context must be considered for an efficient drug design. Besides displaying cell membrane permeability, HDAC6 ZnF-UBP inhibitors should have high affinity in order to compete with abundant ubiquitin substrate inside the cell. Moreover, the versatile interactions that HDAC6 establishes with other partner proteins, such as the Hsp90-HSF1 complex or cortactin could impede an optimal accessibility or conformation for ZnF-UBP domain druggability (36).

In conclusion, the strong impact of a non-functional HDAC6 ZnF-UBP binding domain on several biological processes in MM cells highlights its potential as a drug target. Newly designed ZnF-UBP inhibitors may be of interest to address some of the challenges in MM treatment and to propose an innovative strategy against MM.

## 4. Material and methods

### 4.1. Site-directed mutagenesis, ZnF-UBP^WT^ and ZnF-UBP^RY^ peptide expression and purification

The plasmid encoding for residues 1190 – 1215 of HDAC6 wild-type ZnF-UBP domain (ZnF-UBP^WT^) was obtained from Addgene (Watertown, MA, USA; plasmid #25297). The mutated plasmid encoding for R1155A-Y1156A mutant ZnF-UBP domain (ZnF-UBP^RY^) was generated by site-directed mutagenesis using the wild-type plasmid as DNA template for PCR (forward primer: 5’ – AGGTCTACTGTGGT<u>GC</u>T<u>GC</u>CATCAATGGCCACATGCTCCAACACCATGGA – 3’; reverse primer: 3’ – TCCAGATGACACCA<u>CG</u>A<u>CG</u>GTAGTTACCGGTGTACGAGGTTGTGGTACCT – 5’). PCR products were incubated with 20 U/µL *DpnI* restriction enzyme (ThermoFisher, Waltham, MA, USA) for 1 h to digest parental non-mutated DNA. TOP10 chemically competent *E. coli* cells (ThermoFisher) were transformed, and plasmid DNA was purified with NucleoSpin Plasmid Mini kit (Macherey-Nagel, Düren, Germany) according to the manufacturer instructions. Sanger sequencing of plasmid DNA was performed to verify the nucleotide substitutions (Microsynth, Balgach, Switzerland).

The peptides were then obtained as follows. Wild-type and mutated plasmids were transformed into chemically competent BL21 Star (DE3) *E. coli* cells (Invitrogen, Waltham, MA, USA), which were then grown at 37 °C under agitation (140 rpm) in 2xTY medium supplemented with 50 µg/mL kanamycin until OD_600_ = 0.8. After that, the culture medium was cooled down to 12 °C and 1 mM isopropyl ß-D-1-thiogalactopyranoside (IPTG, Sigma-Aldrich, St. Louis, MO, USA) was added to induce protein expression. After 48 h of incubation, bacteria were harvested by centrifugation at 5000 rpm for 30 min at 4 °C. The supernatant was discarded and the pellet was resuspended in buffer (20 mM Tris-HCl (pH 8), 500 mM NaCl, 20 mM imidazole (pH 8), 10% glycerol, 10 mM β-mercaptoethanol, 1 mM PMSF). Bacterial suspensions were physically lysed by liquid homogenization using a French press (ThermoFisher) and then centrifuged at 15,000 g for 30 min at 4 °C. Lysates were passed through a 0.2 µm filter to remove cell debris and loaded on a HisTrap column (Cytiva, Marlborough, MA, USA) for peptide purification. Purified protein samples were fractionated by FPLC with maximal elution levels at 150 mM of imidazole. The purity of eluted fractions was assessed by 20% SDS-polyacrylamide gel electrophoresis and Coomassie detection.

### 4.2. Fluorescence polarization assay

ZnF-UBP^WT^ and ZnF-UBP^RY^ peptides were serially diluted 1:2 from a starting concentration of 10 µM in a total volume of 60 µL of assay buffer (10 mM HEPES (pH 7.4), 150 mM NaCl) in a 96-well black polystyrene half-area plate (Corning, Corning, NY, USA). Then, the FITC-labeled ubiquitin pentapeptide RLRGG (GenScript, Piscataway, NJ, USA) was added to a final working concentration of 50 nM. Plates were incubated for 5 min under agitation (230 rpm) at RT. Fluorescence polarization was measured on a CLARIOstar plate reader (BMG Labtech, Ortenberg, Germany) at excitation and emission wavelengths of 482 and 530 nm, respectively. When compounds were tested, they were serially diluted 1:2 in assay buffer from a starting concentration of 12.5 µM and wild-type HDAC6 ZnF-UBP domain was added at a working concentration of 2.5 µM, corresponding to approximately 80% of its maximum FP, together with the FITC-labeled ubiquitin pentapeptide RLRGG at 50 nM.

### 4.3. Molecular dynamic simulations

The crystal structure of human HDAC6 zinc finger domain in complex with ubiquitin C-terminal peptide RLRGG (PDB number: 3GV4) served as the template for all molecular dynamic simulations. The proposed R1155A-Y1156A mutation was introduced using the mutagenesis function of Molsoft ICM (http://www.molsoft.com/). Both the wild-type and mutant proteins were simulated in their apo and complexed states for 500 ns. Each system was simulated as triplicates. Parameters for protein atoms were obtained from the standard AMBER FF14SB libraries while customized zinc parameters from the Zinc AMBER Force Field (ZAFF) were employed to accurately represent the zinc ion coordination environment and ion interactions (37, 38). Each system was solvated in a TIP3P water box with a 10 Å buffer from the protein and neutralized with 150 mM K+ and Cl-(39). Systems underwent an initial 500-step steepest descent energy minimization, with constraint applied on the ligand peptide to allow its side chains to relax and form the stabilizing interaction.

Simulations were performed using ACEMD package starting with 5 ns equilibration (40). Systems were then heated from 0 K to 300 K over 100 ps and equilibrated at 300 K and 1 atm for 1 ns using the NPT ensemble. Production runs extended for 500 ns at a timestep of 4 fs employing hydrogen mass partitioning. The coordinates were saved every 100 ps. Long-range electrostatics were computed with Particle Mesh Ewald summation, and the van der Waals interactions were cut off at 9 Å. Analyses focused on conformational changes were conducted using MDANALYSIS and custom scripts (41). The Poisson-Boltzmann electrostatic calculations were carried out using APBS biomolecular solvation software suite version 3.4.1 and the figures were generated using UCSF ChimeraX (42, 43).

### 4.4. CRISPR-Cas9 edition

The human multiple myeloma RPMI 8226 cells (ATCC CCL-155, Manassas, VA, USA) were cultured in complete Roswell Park Memorial Institute (RPMI) 1640 medium (Gibco, Waltham, MA, USA), supplemented with 10% fetal bovine serum (Biowest, Nuaillé, France), 50 U/mL penicillin, 50 µg/mL streptomycin (Gibco), 2 mM L-glutamine (Gibco) at 37 °C in a humified atmosphere with 5% CO_2_.

Knockout HDAC6 (HDAC6^KO^) and ZnF-UBP mutant (HDAC6^RY^) RPMI 8226 cell lines were generated through CRISPR-Cas9 editing. Single-guide RNAs (sgRNA) and the single-stranded DNA donor template (ssODN) for the HDAC6^RY^ RPMI 8226 cells were designed using design tools from Integrated DNA Technologies (IDT, Coralville, IA, USA) (Table 2).

**Table 2.**
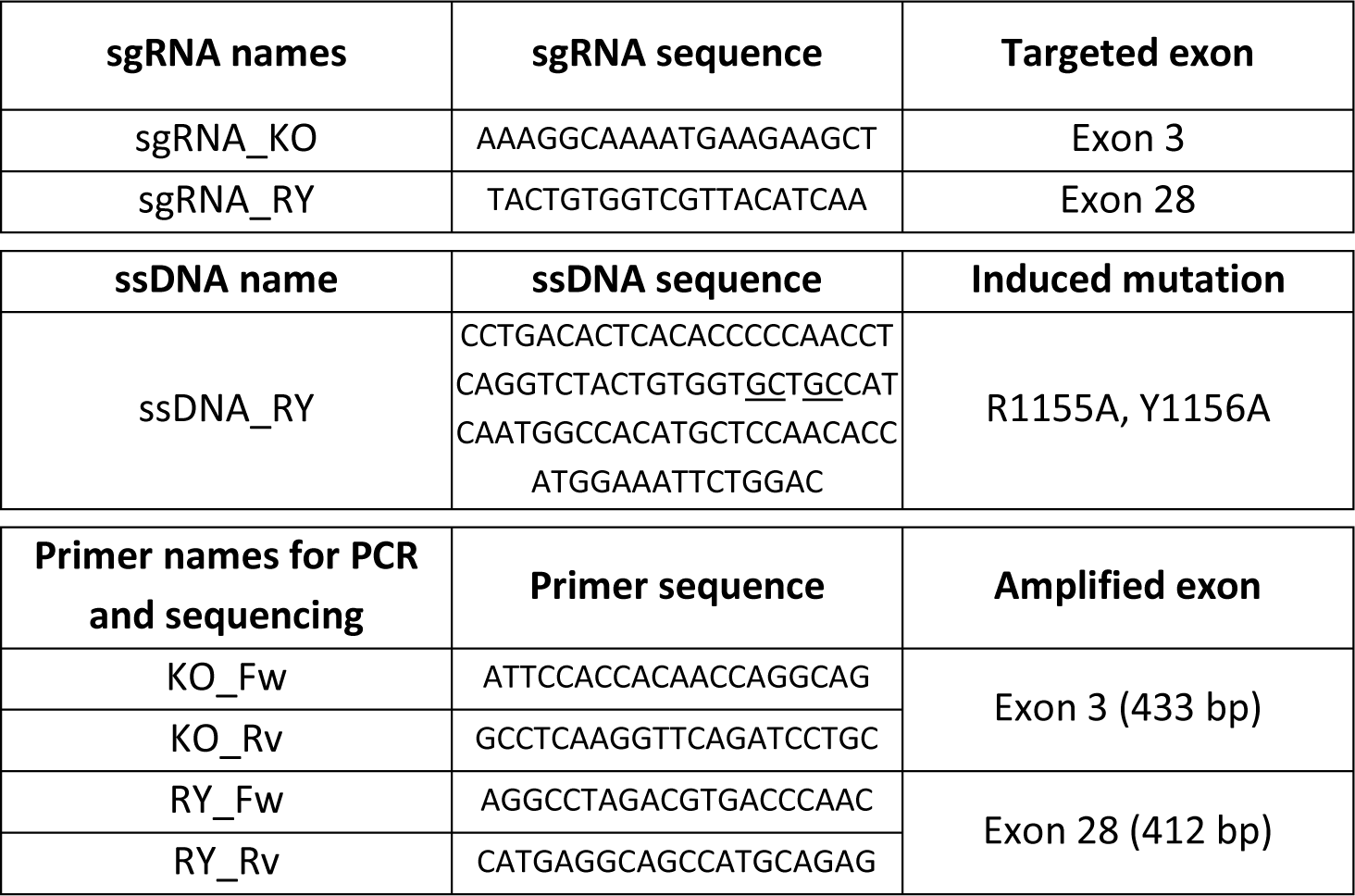
sgRNA, ssDNA and primer sequences (5’ – 3’) for CRISPR-Cas9 edition.

Cas9 ribonucleoprotein complexes (RNP) were assembled by incubating the sgRNA of interest with Alt-R CRISPR-Cas9 tracrRNA ATTO 550 (IDT) at equimolar concentrations at 95 °C for 5 min to ensure hybridization. After cooling down for 10 min at RT, Alt-R HiFi Cas9 Nuclease V3 (IDT) was added and the complex was incubated for 20 min at RT. RPMI 8226 cells (130,000 cells) were washed three times with PBS and resuspended in a final volume of 15 µL of buffer containing the RNA complex (1.8 µM sgRNA, 1.8 µM tracrRNA, 1.5 µM HiFi Cas9) and 1.8 µM electroporation enhancer (IDT). ssODN was also added (3 μM) to obtain the ZnF-UBP mutant cells. Ten µL of this cell suspension were electroporated using the Neon Transfection System (ThermoFisher) with optimized conditions (1150 V, 30 ms, 2 pulses) and immediately diluted in 100 µL of prewarmed RPMI 1640 medium without antibiotics, supplemented with 1 µL HDR enhancer (IDT, to obtain the ZnF-UBP mutant cells). Electroporated cells were then incubated for 24 h at 37 °C, washed three times with PBS and resuspended in sorting buffer (PBS with 1% BSA). Live ATTO 550-positive cells were sorted and individually added to a well containing 100 µL of conditioned medium. Medium was regularly added to the wells to avoid evaporation, and clonal populations were obtained after about 4 weeks of growth. Then, their genomic DNA was extracted using the Wizard Genomic DNA Purification Kit (Promega, Madison, WI, USA), according to the manufacturer instructions. Regions of interest were amplified through polymerase chain reaction (PCR) using designed primers (IDT) (Table 2) and PCR products were sent for Sanger sequencing (Microsynth). For HDAC6^KO^ RPMI 8226 cells, chromatograms were analyzed by the TIDE (Tracking of Indels by DEcomposition) algorithm (http://tide-calculator.nki.nl/, Netherlands Cancer Institute) to obtain the editing efficiency of Cas9.

### 4.5. Western blot

Cells (wild-type, HDAC6^KO^ and HDAC6^RY^ RPMI 8226, 2 × 10^6^) were lysed using NP-40 lysis buffer (ThermoFisher) supplemented with protease inhibitor cocktail (Sigma-Aldrich) and 0.2 M phenylmethylsulfonyl fluoride (ThermoFisher) for 30 min at 4 °C. Lysates were centrifuged at 16,000 g for 10 min at 4 °C and supernatants containing soluble proteins were collected. Proteins were quantified using Coomassie Protein Assay Reagent (ThermoFisher). Protein samples (30 μg) were mixed with loading buffer (50 mM Tris pH 6.8, 6.5% glycerol, 1% sodium dodecyl sulfate, 50 mM dithiothreitol, 0.012% bromophenol blue) and heated to 95 °C for 5 min. Then, proteins were separated in sodium dodecyl sulfate – 8% polyacrylamide gel by electrophoresis at 100 V for 2 h (Bio-Rad, Hercules, CA, USA). Wet transfer to a PVDF membrane (Bio-Rad) was performed in transfer buffer (48 mM Tris, 39 mM glycine, 10% methanol) at 100 V for 90 min. Membranes were blocked in TBS-T (50 mM Tris pH 7.5, 150 mM NaCl, 0.1% Tween 20) containing 5% milk for 1 h and primary antibodies were incubated overnight at 4 °C. The following primary antibodies were used: HDAC6 (Cell Signaling Technology, Danvers, MA, USA, #7558, 1:500), acetylated tubulin (Cell Signaling Technology #3971, 1:1,000), GADPH (Merck, Burlington, MA, USA, MAB374, 1:500). After washing the membranes three times with TBS-T, species-specific HRP-conjugated secondary antibodies (Cell Signaling Technology, all at 1:1,000) were incubated in TBS-T containing 5% milk for 1 h. Proteins were revealed with SuperSignal West Pico PLUS Chemiluminescent Substrate (ThermoFisher) and then imaged using an iBright CL1500 Imaging System (ThermoFisher).

### 4.6. Histone deacetylase catalytic activity

Cells were seeded in a 96-well plate at a density of 15,000 cells per well in 50 µL of medium containing the HDAC substrate N-(4-Methyl-7-aminocoumarinyl)-Nα-(t-butoxycarbonyl)-Nω-acetyllysineamide (21 μM, MAL, Sigma-Aldrich) and incubated for 8 h at 37 °C. Following the incubation, the reaction was stopped by adding 10 µL of 6X RIPA buffer, supplemented with protease inhibitor (Sigma-Aldrich). Then, 160 µL of cold acetonitrile were added and the plate was placed at -80 °C for 10 min. The plate was centrifuged at 5000 g for 10 min at 4 °C, and the supernatants were transferred to a UHPLC-adapted 96-well plate (Nunc 96-well V-bottom polypropylene plate, ThermoFisher) and sealed. The substrate (MAL) and its deacetylated product (dMAL) were measured using a UHPLC-MS method as previously described (44).

### 4.7. Aggresome formation assay

Cells were seeded in a 6-well plate at a density of 3 × 10^5^ cells/mL in 2 mL of medium and assays were performed 48 h later during the log phase of cell growth. Cells were treated with 5 µM MG-132 for 18 h to stimulate aggresome formation. After, they were fixed with 4% paraformaldehyde (Sigma-Aldrich) for 30 min at RT and then permeabilized with a solution of 0.5% Triton X-100 (Sigma-Aldrich) and 3 mM EDTA (pH 8) (Sigma-Aldrich) for 30 min at 4 °C. After the incubation, cells were stained with Aggresome Detection Reagent (1:5000, #ab139486, Abcam, Cambridge, UK) for 30 min in the dark, and then analyzed by flow cytometry (NovoSampler Pro, Agilent, Santa Clara, CA, USA). Aggresome formation levels were detected at excitation and emission wavelengths of 488 and 600 nm, respectively, and mean of fluorescence intensity (MFI) was calculated using FlowJo v10.10 analysis software (Ashland, OR, USA).

### 4.8. Cell growth assay

Cells were seeded in a 96-well plate at a density of 75,000 cells/mL in 200 μL of medium and incubated at 37 °C for 0, 6, 12, 24, 36, 48, 60 or 72 h before adding 50 µL of a solution of 1 mg/mL XTT (ThermoFisher) and continuing the incubation for 4 h. The absorbance was measured at 450 nm using a Cytation 3 plate reader (BioTek, Winooski, VT, USA).

### 4.9. Cell cycle assay

Cells were collected from culture during the log phase of cell growth (500,000 cells), washed once with PBS and fixed in cold 70% ethanol for 30 min at 4 °C. Cells were then washed twice with PBS and incubated with propidium iodide staining solution (2.5 μg/mL PI (BD Biosciences, Franklin Lakes, NJ, USA), 100 µg/mL RNAse A (Promega)) for 2 h at RT. Cells were acquired (NovoSampler Pro) and propidium iodide signal was detected at excitation and emission wavelengths of 488 and 600 nm, respectively. The percentages of cells in G0/G1, S and G2/M phases were obtained using FlowJo v10.10.

### 4.10. Chemical synthesis

Starting building blocks **2a-g** and **3** were purchased at the highest commercial quality from Enamine (Kyiv, Ukraine) and Acros (ThermoFisher), respectively. Solvents for solution phase reactions: N,N’-dimethylformamide (DMF), acetone, acetonitrile (MeCN), methanol (MeOH), ethanol (EtOH) and toluene (PhMe) were purchased in anhydrous grade from Sigma-Aldrich and used without further purification. Ethyl acetate (EtOAc), petroleum ether (PET), cyclohexane, chloroform (CHCl_3_), dichloromethane (CH_2_Cl_2_) and diethyl ether (Et_2_O) were used as received. HPLC-quality acetonitrile (MeCN), HPLC-quality MeOH and MilliQ water were used for HPLC analyses and purifications. Evaporations were conducted under reduced pressure at temperatures less than 40 °C unless otherwise noted. Reactions were monitored by analytical thin layer chromatography (TLC) carried out on ALUGRAM Xtra SIL G/UV254 silica gel plates (Macherey-Nagel). Compounds were visualized with a UV lamp (λ = 254 nm) and/or stained with a solution of potassium permanganate. Determination of purity of compounds was evaluated by analytical HPLC (VWR Hitachi Chromaster, Radnor, PA, USA) on a reversed-phase column (4.6 × 250 mm, 5 µm, 80A, ZORBAX Extend-C18, Agilent), using MeOH/H_2_O or MeCN/H_2_O as the mobile phase (UV detection on a range from 200 to 800 nm). Column chromatography was carried out under positive pressure using 15-40 µm silica gel (Macherey-Nagel) and the indicated solvents. Melting points were recorded in open capillary tubes on a Buchi Melting Point M-560 apparatus (Flawil, Switzerland) and are uncorrected. IR spectra were recorded on an Agilent Cary 630 FT-IR spectrometer. Optical rotations were measured at room temperature on an Applied Biosystems 2720 Thermal Cycler polarimeter (Waltham, MA, USA) using a solution cell of 10 cm optical path. NMR spectra of samples in the indicated solvent were recorded on a Bruker Advance 400 or 500 MHz spectrometer (Billerica, MA, USA) and were calibrated using residual solvent as internal reference. The following abbreviations were used to indicate multiplicities: s = singlet, bs = broad singlet, d = doublet, t = triplet, q = quartet, quint = quintuplet, m = multiplet, and AB = AB quartet. Carbon multiplicities were determined by DEPT-135 experiments. Diagnostic correlations were obtained by two-dimensional COSY, HSQC and HMBC experiments. Low resolution (ESIMS) and/or high-resolution mass spectrometric analyses (HRMS) were obtained by electrospray ionization on a on a Bruker Q-TOF Impact HD apparatus and performed at the PLATeforme INstrumentale d’Analyses (PLATINA) mass spectrometry facility of the Institut de Chimie des Milieux et Matériaux de Poitiers (IC2MP, CNRS-UMR 7285, 86000 Poitiers, France).

#### Compound **5b**

According to the procedure described in the literature (20), compound **4b** (2.1 g, 8.3 mmol, 1 equiv) was suspended in absolute EtOH (80 ml) in a flask equipped with a condenser. A catalytic amount of concentrated H_2_SO_4_ (0.5 ml) was carefully added and the mixture was refluxed until a turbid solution was obtained (17 h). After completion of the reaction (monitored by TLC), the reaction mixture was allowed to cool to 0 °C and was filtered to afford the desired product **5b** as white crystals (1.97 g, 84%). R*_f_* = 0.48 (EtOAc/cyclohexane 50:50); mp 229-231 °C; ^1^H NMR (400 MHz, CDCl_3_) *δ* (ppm): 10.97 (bs, 1H), 8.22 (d, *J* = 2.4 Hz, 1H), 7.68 (dd, *J* = 8.7, 2.4 Hz, 1H), 7.59 (d, *J* = 8.7 Hz, 1H), 4.19 (q, *J* = 7.1 Hz, 2H), 3.06-3.03 (m, 2H), 2.93-2.89 (m, 2H), 1.27 (t, *J* = 7.1 Hz, 3H); ^13^C NMR (100 MHz, CDCl_3_) *δ* (ppm): 173.0, 162.1, 155.1, 147.5, 135.1, 132.4, 128.9, 125.7, 121.9, 61.2, 30.7, 30.2, 14.2; HRMS calcd for C_13_H_14_ClN_2_O_3_ [M+H]^+^ 281.0693, found 281.0686.

#### Compound **6b**

According to the procedure described in the literature (20), K_2_CO_3_ (608 mg, 4.4 mmol, 1.2 equiv) and intermediate **5b** (1.03 g, 3.7 mmol, 1 equiv) were mixed in dry DMF (18.5 ml) under N_2_ atmosphere. Iodomethane (0.45 ml, 7.4 mmol, 2 equiv) was added and the reaction was stirred at RT for 22 h. The resulting mixture was diluted with water (30 ml) and extracted with Et_2_O (4 × 25 ml). The combined organic layers were washed with brine (3 × 30 ml), dried over MgSO_4_, filtered and evaporated. The crude mixture was purified by column chromatography (Et_2_O/PET 10:90 → 45:55) to afford the desired product **6b** as white crystals (1.06 g, 98%). R*_f_* = 0.41 (Et_2_O/PET 60:40); mp 56-58 °C; ^1^H NMR (400 MHz, CDCl_3_) *δ* (ppm): 8.20 (d, *J* = 2.4 Hz, 1H), 7.62 (dd, *J* = 8.7, 2.4 Hz, 1H), 7.51 (d, *J* = 8.7 Hz, 1H), 4.18 (q, *J* = 7.1 Hz, 2H), 3.64 (s, 3H), 3.10 (t, *J* = 6.6 Hz, 2H), 2.92 (t, *J* = 6.7 Hz, 2H), 1.28 (t, *J* = 7.1 Hz, 3H); ^13^C NMR (100 MHz, CDCl_3_) *δ* (ppm): 172.6, 161.2, 155.1, 145.4, 134.3, 132.0, 128.6, 126.0, 121.2, 60.6, 30.1, 30.0, 29.5, 14.2; HRMS calcd for C_14_H_16_ClN_2_O_3_ [M+H]^+^ 295.0849, found 295.0841.

#### Quinazolinone **1b**

According to the procedure described in the literature (20), a solution of the ester **6b** (233 mg, 0.8 mmol, 1 equiv) in a THF/MeOH 1:1 mixture (18 ml), 1 N KOH (3.2 ml, 3.2 mmol, 4 equiv) was added. The mixture was stirred at RT for 22 h. The reaction was diluted with water and the organic solvents were evaporated *in vacuo*. The residue was carefully acidified with aq. 1 N HCl to pH 6-7. The resulting white solid was filtered, washed with water and dried (176 mg, 83%). R*_f_* = 0.10 (Et_2_O); mp 219-221 °C; IR (neat) ν_max_ 1674, 1588, 1413, 1334, 1215, 1139 cm^-1^; ^1^H NMR (400 MHz, DMSO-*d_6_*) *δ* (ppm): 12.23 (bs, 1H), 8.02 (d, *J* = 2.5 Hz, 1H), 7.79 (dd, *J* = 8.7, 2.5 Hz, 1H), 7.57 (d, *J* = 8.7 Hz, 1H), 3.55 (s, 3H), 3.08 (t, *J* = 6.7 Hz, 2H), 2.76 (t, *J* = 6.6 Hz, 2H); ^13^C NMR (100 MHz, DMSO-*d_6_*) *δ* (ppm): 173.8, 160.3, 157.2, 145.3, 134.3, 130.4, 129.0, 125.1, 120.8, 29.9, 29.8, 29.2; HRMS calcd for C_12_H_12_ClN_2_O_3_ [M+H]^+^ 267.0536, found 267.0526; HPLC R_t_ = 19.61 min, purity 98.3%.

#### Compound **5c**

According to the procedure described in the literature (20), compound **4c** (1 g, 4 mmol, 1 equiv) was suspended in absolute EtOH (40 ml) in a flask equipped with a condenser. A catalytic amount of concentrated H_2_SO_4_ (0.2 ml) was carefully added and the mixture was refluxed until a turbid solution was obtained (18 h). After completion of the reaction (monitored by TLC), the reaction mixture was allowed to cool to 0 °C and was filtered to afford the desired product **5c** as white crystals (1.06 g, 95%). R*_f_* = 0.70 (MeOH/CH_2_Cl_2_ 5:95); mp 168-170 °C; ^1^H NMR (400 MHz, DMSO-*d_6_*) *δ* (ppm): 7.59 (d, *J* = 8.8 Hz, 1H), 7.51 (d, *J* = 2.9 Hz, 1H), 7.47 (dd, *J* = 8.8, 3.0 Hz, 1H), 4.06 (q, *J* = 7.1 Hz, 2H), 3.88 (s, 3H), 2.98-2.95 (m, 2H), 2.87-2.84 (m, 2H), 1.15 (t, *J* = 7.1 Hz, 3H); ^13^C NMR (100 MHz, DMSO-*d_6_*) *δ* (ppm): 171.3, 159.8, 158.8, 158.5, 125.1, 123.2, 120.8, 107.0, 60.5, 56.0, 30.2, 27.7, 14.1; HRMS calcd for C_14_H_17_N_2_O_4_ [M+H]^+^ 277.1188, found 277.1181.

#### Compound **6c**

According to the procedure described in the literature (20), K_2_CO_3_ (541 mg, 3.9 mmol, 1.2 equiv) and intermediate **5c** (900 mg, 3.26 mmol, 1 equiv) were mixed in dry DMF (16 ml) under N_2_ atmosphere. Iodomethane (0.4 ml, 6.5 mmol, 2 equiv) was added and the reaction was stirred at RT for 24 h. The resulting mixture was diluted with water (25 ml) and extracted with Et_2_O (4 × 20 ml). The combined organic layers were washed with brine (3 × 25 ml), dried over MgSO_4_, filtered and evaporated. The crude mixture was purified by column chromatography (Et_2_O/PET 20:80 → 75:25) to afford the desired product **6c** as a white microcrystals (542 mg, 57%). R*_f_* = 0.35 (Et_2_O/PET 75:25); mp 125-127 °C; ^1^H NMR (400 MHz, CDCl_3_) *δ* (ppm): 7.60 (d, *J* = 3.0 Hz, 1H), 7.51 (d, *J* = 8.9 Hz, 1H), 7.29 (d, *J* = 8.7 Hz, 1H), 4.18 (q, *J* = 7.1 Hz, 2H), 3.64 (s, 3H), 3.10 (t, *J* = 6.6 Hz, 2H), 2.92 (t, *J* = 6.7 Hz, 2H), 1.28 (t, *J* = 7.1 Hz, 3H); ^13^C NMR (100 MHz, CDCl_3_) *δ* (ppm): 172.7, 162.2, 158.0, 152.4, 141.6, 128.5, 124.3, 120.8, 105.8, 60.5, 55.7, 30.3, 30.0, 29.3, 14.2; HRMS calcd for C_15_H_19_N_2_O_4_ [M+H]^+^ 291.1345, found 291.1342.

#### Quinazolinone **1c**

According to the procedure described in the literature (20), a solution of the ester **6c** (290 mg, 1 mmol, 1 equiv) in a THF/MeOH 1:1 mixture (24 ml), 1 N KOH (4 ml, 4 mmol, 4 equiv) was added. The mixture was stirred at RT for 23 h. The reaction was diluted with water and the organic solvents were evaporated *in vacuo*. The residue was carefully acidified with aq. 1 N HCl to pH 6-7. The resulting white solid was filtered, washed with water and dried (115 mg, 44%). R*_f_* = 0.38 (Et_2_O); mp 204-206 °C; IR (neat) ν_max_ 2934, 1702, 1666, 1491, 1212, 1019, 849, 795 cm^-1^; ^1^H NMR (400 MHz, DMSO-*d_6_*) *δ* (ppm): 12.16 (bs, 1H), 7.51 (d, *J* = 8.9 Hz, 1H), 7.47 (d, *J* = 2.9 Hz, 1H), 7.38 (dd, *J* = 8.9, 3.0 Hz, 1H), 3.86 (s, 3H), 3.56 (s, 3H), 3.06 (t, *J* = 6.7 Hz, 2H), 2.76 (t, *J* = 6.6 Hz, 2H); ^13^C NMR (100 MHz, DMSO-*d_6_*) *δ* (ppm): 173.9, 161.1, 157.4, 154.2, 141.3, 128.5, 123.8, 120.4, 106.0, 55.6, 29.9, 29.7, 28.9; HRMS calcd for C_13_H_15_N_2_O_4_ [M+H]^+^ 263.1032, found 263.1023; HPLC R_t_ = 17.77 min, purity 100%.

#### Compound **5d**

According to the procedure described in the literature (20), compound **4d** (711 g, 3 mmol, 1 equiv) was suspended in absolute EtOH (30 ml) in a flask equipped with a condenser. A catalytic amount of concentrated H_2_SO_4_ (0.2 ml) was carefully added and the mixture was refluxed until a turbid solution was obtained (24 h). After completion of the reaction (monitored by TLC), the reaction mixture was allowed to cool to 0 °C and was filtered to afford the desired product **5d** as white crystals (794 mg, 99%). R*_f_* = 0.82 (MeOH/CH_2_Cl_2_ 10:90); mp 214-215 °C; ^1^H NMR (400 MHz, DMSO-*d_6_*) *δ* (ppm): 12.13 (bs, 1H), 7.86 (s, 1H), 7.57 (dd, *J* = 8.3, 1.8 Hz, 1H), 7.44 (dd, *J* = 8.2 Hz, 1H), 4.04 (q, *J* = 7.1 Hz, 2H), 2.88-2.85 (m, 2H), 2.80-2.76 (m, 2H), 2.41 (s, 3H), 1.14 (t, *J* = 7.1 Hz, 3H); ^13^C NMR (100 MHz, DMSO-*d_6_*) *δ* (ppm): 172.1, 161.6, 155.0, 146.7, 135.6, 135.6, 126.7, 125.1, 120.6, 59.9, 29.8, 28.9, 20.8, 14.1; HRMS calcd for C_14_H_17_N_2_O_3_ [M+H]^+^ 261.1239, found 261.1233.

#### Compound **6d**

According to the procedure described in the literature (20), K_2_CO_3_ (116 mg, 0.8 mmol, 1.2 equiv) and intermediate **5d** (182 mg, 0.7 mmol, 1 equiv) were mixed in dry DMF (4 ml) under N_2_ atmosphere. Iodomethane (0.1 ml, 1.4 mmol, 2 equiv) was added and the reaction was stirred at RT for 19 h. The resulting mixture was diluted with water (10 ml) and extracted with Et_2_O (4 × 10 ml). The combined organic layers were washed with brine (2 × 15 ml), dried over MgSO_4_, filtered and evaporated. The crude mixture was purified by column chromatography (Et_2_O/PET 50:50 → 90:10) to afford the desired product **6d** as a white solid (157 mg, 82%). R*_f_* = 0.46 (Et_2_O); mp 97-99 °C; ^1^H NMR (400 MHz, CDCl_3_) *δ* (ppm): 8.03 (s, 1H), 7.51 (dd, *J* = 8.4, 2.0 Hz, 1H), 7.47 (d, *J* = 8.2 Hz, 1H), 4.18 (q, *J* = 7.1 Hz, 2H), 3.63 (s, 3H), 3.10 (t, *J* = 6.8 Hz, 2H), 2.92 (t, *J* = 6.7 Hz, 2H), 2.46 (s, 3H), 1.27 (t, *J* = 7.1 Hz, 3H); ^13^C NMR (100 MHz, CDCl_3_) *δ* (ppm): 172.7, 162.3, 153.8, 144.9, 136.4, 135.4, 126.7, 126.0, 119.9, 60.5, 30.3, 29.9, 29.4, 21.2, 14.2; HRMS calcd for C_15_H_19_N_2_O_3_ [M+H]^+^ 275.1396, found 275.1391.

#### Quinazolinone **1d**

According to the procedure described in the literature (20), a solution of the ester **6d** (100 mg, 0.36 mmol, 1 equiv) in a THF/MeOH 1:1 mixture (4 ml), 2 N KOH (0.7 ml, 1.4 mmol, 4 equiv) was added. The mixture was stirred at RT for 20 h. The reaction was diluted with water and the organic solvents were evaporated *in vacuo*. The residue was carefully acidified with aq. 1 N HCl to pH 6-7. The resulting white solid was filtered, washed with water and dried (82 mg, 91%). R*_f_* = 0.22 (Et_2_O); mp 207-209 °C; IR (neat) ν_max_ 2575, 1709, 1650, 1567, 1406, 1037, 830, 782 cm^-1^; ^1^H NMR (400 MHz, DMSO-*d_6_*) *δ* (ppm): 7.93 (bs, 1H), 7.69 (dd, *J* = 8.4, 1.8 Hz, 1H), 7.63 (d, *J* = 8.3 Hz, 1H), 3.57 (s, 3H), 3.19 (t, *J* = 7.0 Hz, 2H), 2.81 (t, *J* = 7.1 Hz, 2H), 2.45 (s, 3H); ^13^C NMR (125 MHz, DMSO-*d_6_*) *δ* (ppm): 173.1, 160.2, 159.1, 139.8, 137.7, 136.6, 126.0, 122.8, 118.7, 30.7, 30.2, 28.2, 20.9; HRMS calcd for C_13_H_15_N_2_O_3_ [M+H]^+^ 247.1083, found 247.1076; HPLC R_t_ = 18.45 min, purity 100%.

#### Compound **5e**

According to the procedure described in the literature (20), compound **4e** (2.1 g, 8.3 mmol, 1 equiv) was suspended in absolute EtOH (80 ml) in a flask equipped with a condenser. A catalytic amount of concentrated H_2_SO_4_ (0.5 ml) was carefully added and the mixture was refluxed until a turbid solution was obtained (19 h). After completion of the reaction (monitored by TLC), the reaction mixture was allowed to cool to 0 °C and was filtered to afford the desired product **5e** as yellow crystals (1.72 g, 74%). R*_f_* = 0.41 (EtOAc/cyclohexane 50:50); mp 164-166 °C; ^1^H NMR (400 MHz, DMSO-*d_6_*) *δ* (ppm): 12.37 (bs, 1H), 8.04 (d, *J* = 8.5 Hz, 1H), 7.55 (dd, *J* = 1.7 Hz, 1H), 7.46 (dd, *J* = 8.5, 1.9 Hz, 1H), 4.04 (q, *J* = 7.1 Hz, 2H), 2.88 (t, *J* = 6.8 Hz, 2H), 2.79 (t, *J* = 6.7 Hz, 2H), 1.14 (t, *J* = 7.1 Hz, 3H); ^13^C NMR (100 MHz, DMSO-*d_6_*) *δ* (ppm): 172.0, 161.1, 157.8, 149.7, 139.0, 127.9, 126.4, 125.9, 119.7, 60.0, 29.7, 29.1, 14.1; HRMS calcd for C_13_H_14_ClN_2_O_3_ [M+H]^+^ 281.0693, found 281.0689.

#### Compound **6e**

According to the procedure described in the literature (20), K_2_CO_3_ (829 mg, 6 mmol, 1.2 equiv) and intermediate **5e** (1.4 g, 5 mmol, 1 equiv) were mixed in dry DMF (18.5 ml) under N_2_ atmosphere. Iodomethane (0.45 ml, 7.4 mmol, 2 equiv) was added and the reaction was stirred at RT for 16 h. The resulting mixture was diluted with water (35 ml) and extracted with Et_2_O (4 × 30 ml). The combined organic layers were washed with brine (3 × 40 ml), dried over MgSO_4_, filtered and evaporated. The crude mixture was purified by column chromatography (Et_2_O/PET 10:90 → 60:40) to afford the desired product **6e** as white crystals (1.37 g, 93%). R*_f_* = 0.35 (Et_2_O/PET 60:40); mp 115-116 °C; ^1^H NMR (400 MHz, CDCl_3_) *δ* (ppm): 8.17 (d, *J* = 8.6 Hz, 1H), 7.57 (dd, *J* = 2.0 Hz, 1H), 7.37 (d, *J* = 8.6, 2.0 Hz, 1H), 4.19 (q, *J* = 7.1 Hz, 2H), 3.63 (s, 3H), 3.10 (t, *J* = 6.4 Hz, 2H), 2.92 (t, *J* = 6.7 Hz, 2H), 1.29 (t, *J* = 7.1 Hz, 3H); ^13^C NMR (100 MHz, CDCl_3_) *δ* (ppm): 172.6, 161.6, 156.2, 147.8, 140.0, 128.2, 126.9, 126.4, 118.6, 60.6, 30.0, 29.9, 29.5, 14.2; HRMS calcd for C_14_H_16_ClN_2_O_3_ [M+H]^+^ 295.0849, found 295.0850.

#### Quinazolinone **1e**

According to the procedure described in the literature (20), a solution of the ester **6e** (1.19 g, 4 mmol, 1 equiv) in a THF/MeOH 1:1 mixture (48 ml), 2 N KOH (8 ml, 16 mmol, 4 equiv) was added. The mixture was stirred at RT for 17 h. The reaction was diluted with water and the organic solvents were evaporated *in vacuo*. The residue was carefully acidified with aq. 1 N HCl to pH 6-7. The resulting white solid was filtered, washed with water and dried (1.08 g, quant.). R*_f_* = 0.10 (Et_2_O); mp 250-252 °C; IR (neat) ν_max_ 1722, 1640, 1590, 1419, 1174, 872 cm^-1^; ^1^H NMR (500 MHz, DMSO-*d_6_*) *δ* (ppm): 12.24 (bs, 1H), 8.05 (d, *J* = 8.5 Hz, 1H), 7.54 (d, *J* = 1.9 Hz, 1H), 7.47 (dd, *J* = 8.5, 1.9 Hz, 1H), 3.53 (s, 3H), 3.07 (t, *J* = 6.7 Hz, 2H), 2.76 (t, *J* = 6.7 Hz, 2H); ^13^C NMR (100 MHz, DMSO-*d_6_*) *δ* (ppm): 173.7, 160.7, 158.3, 147.6, 138.8, 128.3, 126.5, 125.7, 118.4, 29.7, 29.7, 29.2; HRMS calcd for C_12_H_12_ClN_2_O_3_ [M+H]^+^ 267.0536, found 267.0531; HPLC R_t_ = 19.63 min, purity 100%.

#### Compound **5f**

According to the procedure described in the literature (20), compound **4f** (395 mg, 1.5 mmol, 1 equiv) was suspended in absolute EtOH (14 ml) in a flask equipped with a condenser. A catalytic amount of concentrated H_2_SO_4_ (0.1 ml) was carefully added and the mixture was refluxed until a turbid solution was obtained (17 h). After completion of the reaction (monitored by TLC), the reaction mixture was allowed to cool to 0 °C and was filtered to afford the crude. The resulting solid was purified by column chromatography with EtOAc as eluent to yield compound **5f** as a white powder (354 mg, 81%) R*_f_* = 0.60 (Et_2_O); mp 165-167 °C; ^1^H NMR (400 MHz, DMSO-*d_6_*) *δ* (ppm): 12.67 (bs, 1H), 8.31 (d, *J* = 8.7 Hz, 1H), 8.25 (dd, *J* = 2.1 Hz, 1H), 8.19 (dd, *J* = 8.7, 2.2 Hz, 1H), 4.07 (q, *J* = 7.1 Hz, 2H), 2.95 (t, *J* = 6.6 Hz, 2H), 2.84 (t, *J* = 6.5 Hz, 2H), 1.17 (t, *J* = 7.1 Hz, 3H); ^13^C NMR (100 MHz, DMSO-*d_6_*) *δ* (ppm): 172.0, 160.6, 158.7, 151.1, 148.9, 128.0, 125.2, 121.6, 119.7, 60.0, 29.6, 29.0, 14.1; HRMS calcd for C_13_H_14_N_3_O_5_ [M+H]^+^ 292.0933, found 292.0930.

#### Compound **6f**

According to the procedure described in the literature (20), K_2_CO_3_ (152 mg, 1.1 mmol, 1.2 equiv) and intermediate **5f** (270 mg, 0.9 mmol, 1 equiv) were mixed in dry DMF (5 ml) under N_2_ atmosphere. Iodomethane (0.11 ml, 1.8 mmol, 2 equiv) was added and the reaction was stirred at RT for 17 h. The resulting mixture was diluted with water (15 ml) and extracted with Et_2_O (4 × 30 ml). The combined organic layers were washed with brine (3 × 20 ml), dried over MgSO_4_, filtered and evaporated. The crude mixture was purified by column chromatography (Et_2_O/PET 10:90 → 75:25) to afford the desired product **6f** as yellow crystals (171 mg, 60%). R*_f_* = 0.30 (Et_2_O/PET 75:25); mp 119-121 °C; ^1^H NMR (400 MHz, CDCl_3_) *δ* (ppm): 8.40 (s, 1H), 8.39 (d, *J* = 11.7 Hz, 1H), 8.17 (dd, *J* = 8.8, 2.2 Hz, 1H), 4.19 (q, *J* = 7.1 Hz, 2H), 3.67 (s, 3H), 3.15 (t, *J* = 6.2 Hz, 2H), 2.96 (t, *J* = 6.7 Hz, 2H), 1.30 (t, *J* = 7.1 Hz, 3H); ^13^C NMR (100 MHz, CDCl_3_) *δ* (ppm): 172.6, 161.1, 157.4, 151.4, 147.3, 128.7, 124.3, 122.7, 120.0, 60.8, 30.3, 29.9, 29.6, 14.3; HRMS calcd for C_14_H_16_N_3_O_5_ [M+H]^+^ 306.1090, found 306.1088.

#### Quinazolinone **1f**

According to the procedure described in the literature (20), a solution of the ester **6f** (100 mg, 0.36 mmol, 1 equiv) in a THF/MeOH 1:1 mixture (4 ml), 2 N KOH (0.7 ml, 1.4 mmol, 4 equiv) was added. The mixture was stirred at RT for 20 h. The reaction was diluted with water and the organic solvents were evaporated *in vacuo*. The residue was carefully acidified with aq. 1 N HCl to pH 6-7. The resulting yellow solid was filtered, washed with water and dried (103 mg, 89%). R*_f_* = 0.17 (Et_2_O); mp 225-227 °C; IR (neat) ν_max_ 3093, 1721, 1650, 1594, 1527, 1340, 1174, 884 cm^-1^; ^1^H NMR (400 MHz, DMSO-*d_6_*) *δ* (ppm): 12.25 (bs, 1H), 8.33 (d, *J* = 8.6 Hz, 1H), 8.23 (d, *J* = 2.0 Hz, 1H), 8.20 (dd, *J* = 8.6, 2.1 Hz, 1H), 3.59 (s, 3H), 3.14 (t, *J* = 6.6 Hz, 2H), 2.79 (t, *J* = 6.5 Hz, 2H); ^13^C NMR (100 MHz, DMSO-*d_6_*) *δ* (ppm): 173.7, 160.3, 159.2, 151.0, 146.8, 128.6, 123.8, 121.6, 119.8, 30.0, 29.7, 29.3; HRMS calcd for C_12_H_12_N_3_O_5_ [M+H]^+^ 278.0777, found 278.0776; HPLC R_t_ = 18.83 min, purity 98.0%.

#### Compound **5g**

According to the procedure described in the literature (20), compound **4g** (1.86 g, 8 mmol, 1 equiv) was suspended in absolute EtOH (80 ml) in a flask equipped with a condenser. A catalytic amount of concentrated H_2_SO_4_ (0.5 ml) was carefully added and the mixture was refluxed until a turbid solution was obtained (22 h). After completion of the reaction (monitored by TLC), the reaction mixture was diluted with water and extracted with EtOAc (3 × 50 ml). The combined organic layers were washed with brine (3 × 100 ml), dried over MgSO_4_, filtered and evaporated. The crude mixture was purified by column chromatography using EtOAc as eluent to afford the desired product **5g** a white powder (1.16 g, 56%). R*_f_* = 0.84 (MeOH/CH_2_Cl_2_ 10:90); mp 184-186 °C; ^1^H NMR (400 MHz, CDCl_3_) *δ* (ppm): 12.06 (bs, 1H), 8.14 (d, *J* = 8.1 Hz, 1H), 7.44 (s, 1H), 7.25 (dd, *J* = 8.0, 1.1 Hz, 1H), 4.16 (q, *J* = 7.2 Hz, 2H), 3.10 (t, *J* = 7.1 Hz, 2H), 2.93 (t, *J* = 7.1 Hz, 2H), 2.47 (s, 3H), 1.24 (t, *J* = 7.1 Hz, 3H); ^13^C NMR (100 MHz, CDCl_3_) *δ* (ppm): 172.6, 163.9, 154.9, 149.3, 145.7, 128.0, 127.0, 126.1, 118.2, 60.8, 30.7, 30.0, 21.9, 14.1; HRMS calcd for C_14_H_17_N_2_O_3_ [M+H]^+^ 261.1239, found 261.1237.

#### Compound **6g**

According to the procedure described in the literature (20), K_2_CO_3_ (664 mg, 4.8 mmol, 1.2 equiv) and intermediate **5g** (1.05 g, 4 mmol, 1 equiv) were mixed in dry DMF (20 ml) under N_2_ atmosphere. Iodomethane (0.5 ml, 8 mmol, 2 equiv) was added and the reaction was stirred at RT for 16 h. The resulting mixture was diluted with water (100 ml) and extracted with Et_2_O (4 × 20 ml). The combined organic layers were washed with brine (2 × 50 ml), dried over MgSO_4_, filtered and evaporated. The crude mixture was purified by column chromatography (Et_2_O/PET 50:50 → 90:10) to afford the desired product **6g** as white microcrystals (1.0 g, 91%). R*_f_* = 0.56 (Et_2_O); mp 114-116 °C; ^1^H NMR (400 MHz, CDCl_3_) *δ* (ppm): 8.12 (d, *J* = 8.1 Hz, 1H), 7.37 (s, 1H), 7.24 (dd, *J* = 8.3, 1.1 Hz, 1H), 4.18 (q, *J* = 7.1 Hz, 2H), 3.63 (s, 3H), 3.10 (t, *J* = 6.8 Hz, 2H), 2.92 (t, *J* = 6.8 Hz, 2H), 2.47 (s, 3H), 1.28 (t, *J* = 7.1 Hz, 3H); ^13^C NMR (100 MHz, CDCl_3_) *δ* (ppm): 172.7, 162.2, 154.7, 147.0, 144.8, 127.9, 126.6, 126.4, 117.8, 60.5, 30.3, 29.8, 29.5, 21.7, 14.2; HRMS calcd for C_15_H_19_N_2_O_3_ [M+H]^+^ 275.1396, found 275.1392.

#### Quinazolinone **1g**

According to the procedure described in the literature (20), a solution of the ester **6g** (912 mg, 3.6 mmol, 1 equiv) in a THF/MeOH 1:1 mixture (42 ml), 2 N KOH (7 ml, 14 mmol, 4 equiv) was added. The mixture was stirred at RT for 4 h. The reaction was diluted with water and the organic solvents were evaporated *in vacuo*. The residue was carefully acidified with aq. 1 N HCl to pH 6-7. The resulting white solid was filtered, washed with water and dried (699 mg, 85%). R*_f_* = 0.11 (Et_2_O); mp 256-258 °C; IR (neat) ν_max_ 2991, 1724, 1640, 1619, 1592, 1420, 1181, 785 cm^-1^; ^1^H NMR (400 MHz, DMSO-*d_6_*) *δ* (ppm): 12.18 (bs, 1H), 7.98 (d, *J* = 8.1 Hz, 1H), 7.36 (s, 1H), 7.29 (d, *J* = 8.1 Hz, 1H), 3.54 (s, 3H), 3.07 (t, *J* = 6.6 Hz, 2H), 2.76 (t, *J* = 6.6 Hz, 2H), 2.44 (s, 3H); ^13^C NMR (125 MHz, DMSO-*d_6_*) *δ* (ppm): 173.8, 161.2, 156.5, 146.7, 144.6, 127.7, 126.3, 126.1, 117.3, 29.8, 29.5, 29.0, 21.3; HRMS calcd for C_13_H_15_N_2_O_3_ [M+H]^+^ 247.1083, found 247.1081; HPLC R_t_ = 18.01 min, purity 97.8%.

### 4.11. Molecular docking

The structure of the HDAC6 ZnF-UBP domain in complex with inhibitors (PDB number: 6CE6) was downloaded from Protein Data Bank (20). The structures of compounds **1a-g** were sketched in the ligedit module implemented in the MolSoft ICM-Pro package (http://www.molsoft.com/). The ZnF-UBP binding site was defined based on the ligand present in the reference PDB (Fig. 6) and enclosed 6 Å cubic grid. The compounds were then docked using a thoroughness value of 10. A maximum of 10 docked conformations per compound was generated. A radial convolutional neural net (RTCNN) score was used to rank the docked compounds (45). The RTCNN score is an abstract value that employs trained features to separate binders from non-binders.

### 4.12. RNA isolation and RNA-seq analysis

RNA was isolated from each cell line using Aurum total RNA mini kit (Bio-Rad) and following manufacturer instructions. In brief, 2 x 10^6^ cells of each line were washed in PBS and then resuspended in 350 μL lysis solution (supplemented with 1% β-mercaptoethanol (Sigma-Aldrich)). 350 μL of 70% ethanol were added and the mixture was transferred to a RNA binding column. After centrifugation, the column was washed with 700 μL of wash solution and 80 μL of DNAse I solution were added to digest DNA. The column was washed again twice and RNA was eluted by adding 50 μL of elution buffer. The RNA quality control of all samples was checked prior to sequencing (RNA integrity number: 10). Sequencing was performed by an Illumina NovaSeq 6000 System using the library TruSeq HT stranded (100 bp single-end reads). After sequencing quality control (FastQC), the reads were aligned to the GRCh38 Ensembl human genome reference with STAR v.2.7.4 (46). The aligned reads were assessed for gene features with HTSeq v0.9.1 and the matrix of counts per genes was produced. The differential expression analysis was performed with the statistical analysis R/Bioconductor package *edgeR* 1.34.1 (47). After normalization, the poorly detected genes were filtered out and out of 60,666 total genes, 21,394 genes with a count above 10 were kept. Among them, only those genes with a fold change 2 threshold and a p-value < 0.05 (with a false discovery rate of 5%) were considered as differentially expressed. Gene ontology enrichment analysis was performed through the R package *clusterProfiler* (48). The RNA-Seq data have been deposited in the NCBI Gene Expression Omnibus with GEO Series accession GSE263138.

### 4.13. IL-6 and IL-10 production measurement

Cells were seeded in a 96-well plate at a density of 5 × 10^5^ cells/mL in 200 μL of medium with 100 ng/mL LPS (InvivoGen, San Diego, CA, USA) and incubated at 37 °C for 24 h. Then, the plate was centrifuged at 200 g for 2 min at RT and supernatants were transferred to another plate for IL-6 and IL-10 quantification. The concentrations of the cytokines were measured by ELISA using two commercially available kits from BioLegend (#430501 for IL-6 and #430601 for IL-10, San Diego, CA, USA). The detection limits for IL-6 and IL-10 were 7.8 pg/mL and 3.9 pg/mL, respectively.

### 4.14. Statistical analysis

Data representation and statistical analyses were performed using GraphPad Prism 9 (GraphPad, La Jolla, CA, USA). Sample distribution was tested for normality (Shapiro-Wilk, α = 0.05) prior to group comparison. The details of the tests used and levels of significance are mentioned in the figure legends.

## Author contributions

Conceptualization, PB and MC; methodology, RR, IFC, AR, FA, MU, SH, CB, PB and MC; biological evaluation, RR, IFC and MU; synthesis of compounds, AR and PB; writing—original draft preparation, RR and MC; writing—review and editing, RR, SH, CB, PB and MC; supervision, PC and MC; project administration, MC. All authors have read and approved the final manuscript.

## Acknowledgements

We thank Aurélie Gouiller and Olivier Petermann for their technical support and La Ligue Contre le Cancer (Comité Charente-Maritime) for their financial support.

